# A Structural Principle for Macroscopic Neural Dynamics Correlations

**DOI:** 10.64898/2026.06.14.729168

**Authors:** Qihang Wu, Quan Wen, Cirong Liu

## Abstract

A central question in neuroscience is how the brain’s structural connectivity gives rise to its emergent, correlated dynamics. These large-scale dynamical correlations underlie functional networks that support cognitive functions. Here, we identify coupling correlation—the similarity between the input connectivity profiles of brain regions—as a key structural determinant of macroscopic neural dynamical correlation. Using dynamical mean-field theory (DMFT) and numerical simulations of random neural network models, we demonstrate that coupling correlation quantitatively governs dynamical correlation. The functional form of this structure–function mapping is dictated by the eigenvalue spectrum of the coupling correlation matrix: networks with bulk eigenspectra exhibit an exact linear relationship, whereas biologically plausible long-tailed spectra yield an approximately linear mapping except when the magnitude of coupling correlation approaches unity. Particularly, a long-tailed spectrum is necessary to reproduce the appropriate magnitude and size-invariance of coupling correlations observed in empirical data, thereby sustaining non-vanishing dynamical correlations that may support brain function in large systems. The theoretical prediction of approximate linearity is consistently validated using empirical datasets that include both structural coupling and neural dynamics in humans, mice, and *Drosophila*. Together, these results provide a mechanistic and quantitative framework linking macroscopic brain network structure to emergent neural dynamics—an essential step toward a theory of structure–function relationship in the brain.

**Significance Statement:** How the brain’s wiring gives rise to its coordinated activity is a fundamental unsolved problem in neuroscience. Prior work has identified correlations between structural and functional connectivity, but these relationships lacked a mechanistic, first-principles explanation. Here, we derive an analytical framework using Dynamical Mean-Field Theory and random neural network models to show that a single structural statistic—*coupling correlation*, the similarity between the input connectivity profiles of brain regions—linearly and causally determines the magnitude of correlated neural dynamics. We further show that a long-tailed eigenvalue spectrum in biological structural connectivity is necessary to sustain the strong, size-invariant functional correlations observed across species. Validated in humans, mice, and *Drosophila* using multiple imaging and connectome modalities, this principle may provide a quantitative bridge between structural connectomics and emergent brain dynamics, with implications extending to a broad class of complex networked systems.

## 1 Main

Large-scale neural dynamics exhibit rich spatiotemporal structure but remain difficult to interpret due to high dimensionality and chaos. A common strategy is to analyze pairwise dynamical correlations between regional activities [1–3]. These correlations reveal modular organization [4, 5] and form functional networks that support cognition, with their disruption marking neurological disorders [6–8].

Despite extensive empirical characterization, the structural origin of these correlations remains unresolved. Early work attributed them to direct anatomical coupling [9–13], but strong correlations frequently arise between regions without direct links [14]. To address this, network communication measures—such as shortest path length, search information, and communicability—have been proposed [15–17]. Linear regression using these measures can achieve statistically significant correlations with dynamical correlation (maximum *r* ≈ 0.4–0.6). While these findings demonstrate that structure constrains dynamics, the relationship is correlational rather than causal: the measures are heuristically designed rather than derived from network model, and the linear form of the regression lacks a theoretical foundation. Moreover, some approaches apply ad hoc transformations (e.g., logarithmic or inverse transformations) without principled justification.

A key limitation of these approaches is their focus on pairwise interaction or communication. In contrast, correlations may arise because two regions are embedded similarly within the global network, rather than because they are efficiently connected [18]. Capturing this mechanism requires a global, rather than path-based, description of network structure [19, 20]. Finally, establishing a causal relationship requires a model that considers full network interactions rather than just pairwise paths.

Random neural networks provide a tractable framework for this problem. Unlike previous analytical works that assumed linear or feedforward architectures [18, 21], they naturally incorporate recurrent nonlinear dynamics similar to biological brains, while remaining analytically accessible via dynamical mean-field theory (DMFT) [22– 25]. However, conventional models assume independent couplings scaling as 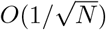, leading to vanishing correlations of the same order—far smaller than those observed in biological systems.

Here, we introduce coupling correlation, defined as the Pearson correlation between the incoming coupling vectors of two units. This quantity captures the similarity of their network-wide inputs, shifting the focus from point-to-point connectivity to global structural embedding. We show that, within a DMFT framework, despite strong nonlinearities in single-unit dynamics, coupling correlation emerges as the exact linear determinant of dynamical correlations at the disorder-average level. While coupling correlation is closely related to empirically used measures such as the matching index [19, 20], these quantities can be understood as heuristic approximations of coupling correlation, a more general structural statistic, whose relationship with dynamical correlation is derived from first principles in this study. Furthermore, we demonstrate that the spectral properties of the coupling correlation matrix govern the validity of this linear mapping: simulated networks with bulk spectra exhibit exact linearity, whereas real brain networks—including *Drosophila*, mouse, and human—display long-tailed spectra, yielding approximate linearity that goes nonlinear only as coupling correlation approaches unity. Finally, to connect the disorder-average prediction of DMFT with single-instance empirical data, we perform a binning analysis that estimates the expectation of dynamical correlation as a function of coupling correlation. Across different experimental data, this binned relationship is strongly linear(*R*^2^ *>* 0.9), confirming the theoretical prediction at the ensemble level.

## 2 Random neural network models and the DMFT equation

We begin by introducing the random neural network models and the theoretical framework of dynamical mean-field theory (DMFT). A random neural network consists of units coupled through weighted directed connections that specify the strength of interactions (**Fig.1a**). For a network with *N* units, the dynamics of the *i*-th unit are given by

**Fig. 1.**
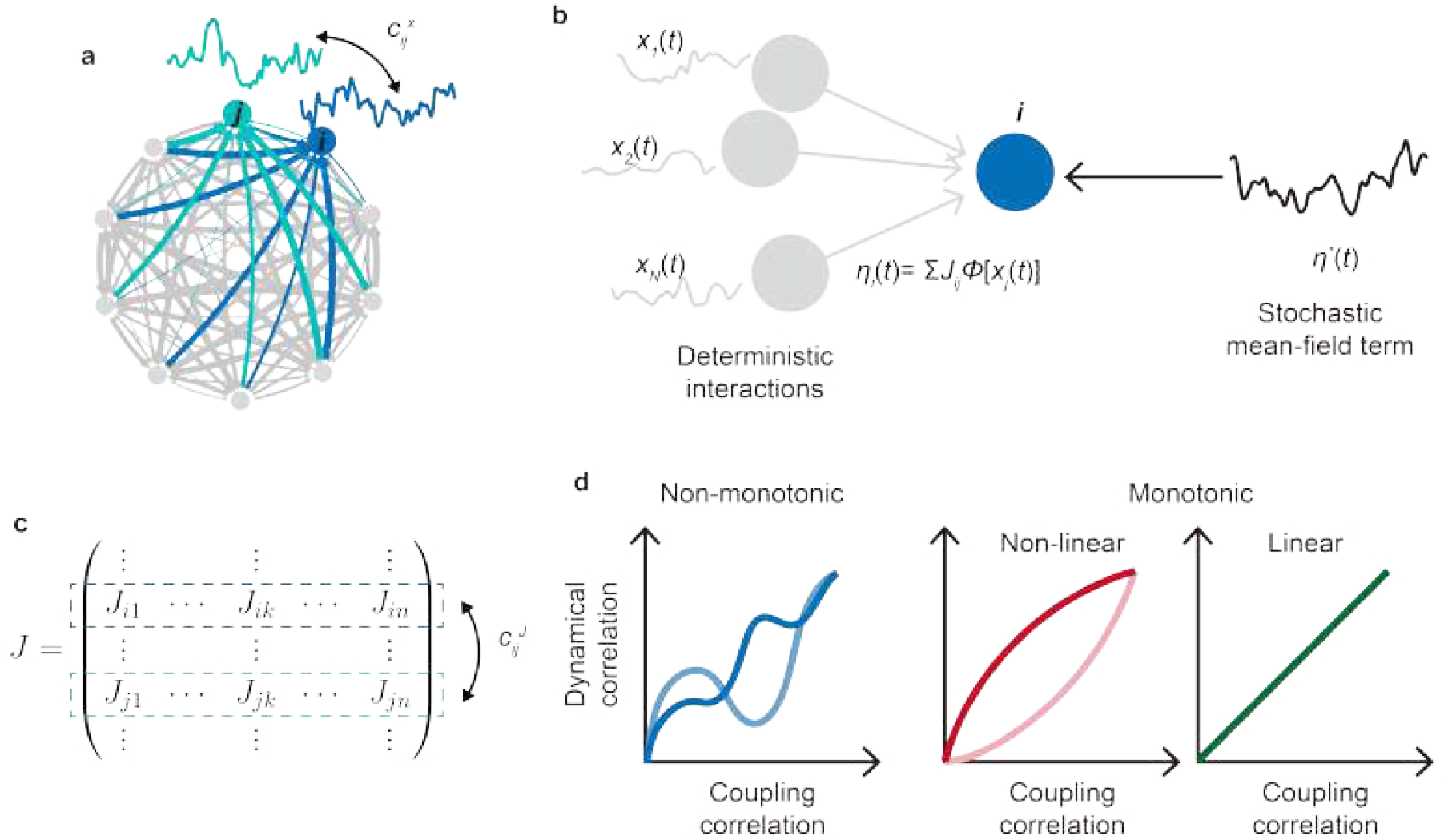
Structure and dynamics of random neural networks. **a**, In the network, all the units are recurrently connected. Units *i* (blue) and *j* (green) each receive inputs from a large population of other units, whose collective influence governs their dynamics and dynamical correlation 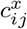. **b**, Schematic of the dynamical mean-field theory (DMFT). In the full network, without external drive, the collective interactions from other units form a deterministic drive *η*_*i*_(*t*) to unit *i*, which replaced by an effective stochastic mean-field term *η*^∗^(*t*) in the DMFT formulation. **c**, The couplings are represented by he matrix **J**. The similarity between the incoming couplings to units *i* (blue arrows in **a** and the blue dashed box here) and (green arrows in **a** and the green dashed box here)—quantified by the coupling correlation 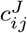, which is defined as the correlation coefficient of the *i*th and *j*th rows of **J. d**, The dependence of dynamical correlation on coupling correlation is expected to be monotonic (middle and right), rather than non-monotonic (left), but can be either linear (right) or nonlinear middle), which calls for theoretical analysis to determine.

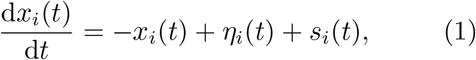

where 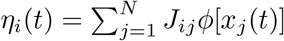 denotes the collective recurrent input from all other units, and *s*_*i*_ represents an external drive. *J*_*ij*_ is the coupling from unit *j* to unit *i*, and *ϕ*(·) is a nonlinear activation function. Conventionally, 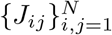 are independent and identically distributed (i.i.d.) Gaussian random variables with a zero mean, *µ*_*J*_ = E[*J*_*ij*_] = 0 (ensuring excitatory–inhibitory balance), and a variance 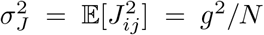. When modeling macroscopic biological neural systems, each unit corresponds to a local brain region comprising large populations of neurons, with *x*_*i*_(*t*) representing the macroscopic neural activity of that region, and *J*_*ij*_ encoding the anatomical connection strength from unit *i* to *j*. The pairwise dynamical correlation 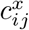 is defined as the correlation coefficient between the dynamics *x*_*i*_ and *x*_*j*_ (**Fig.1a**), corresponding to what is typically referred to as functional connectivity in neuroscience. While anatomical connectivity in the brain is fixed, the random couplings in the model offer a more tractable framework for theoretical analysis. By preserving key features—such as the coupling strength parameter *g*—and randomizing irrelevant details, they enable the identification of structural determinants that causally shape macroscopic dynamical properties, like chaos and dynamical correlation. Predictions derived from these networks can then be directly compared with empirical neural data, bridging theory and experiments.

The rationale of DMFT can be best illustrated in the absence of an external drive. In this case, directly solving the original system with *N* coupled deterministic differential equations is intractable when *N* is large. The DMFT provides a principled reduction (**Fig.1b**): in the thermodynamic limit (*N* → ∞), the collective recurrent input *η*_*i*_(*t*) is proved to converge to a one-dimensional zero-centered Gaussian process *η*^∗^(*t*) with autocorrelation ⟨*η*^∗^(*t*)*η*^∗^(*t*′)⟩_*t*_ = *g*^2^ ∑_*i*_⟨*ϕ*_*i*_(*t*)*ϕ*_*i*_(*t*′)⟩*/N* where ⟨·⟩_*t*_ and ⟨·⟩ represent temporal and dynamical trajectory averages, respectively. Importantly, the autocorrelation magnitude is constrained by the structural coupling statistics *g*, illustrating a causal relationship between structure and dynamics. As a result, the high-dimensional deterministic system is replaced by an effective one-dimensional stochastic differential equation for a representative unit, known as the DMFT equation. Solving this effective equation under the self-consistency condition—i.e. ensuring that the statistics of *η*^∗^(*t*) match those generated by the dynamics in equation (1), yields the macroscopic statistical properties of the original network, including its spatiotemporal correlation structure.

Here, unlike previous works on fully random networks, we consider the scenario in which the incoming structural couplings of different units are correlated (**Fig.1c**). This is captured by the simplified construction

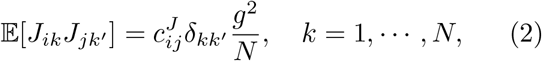

where 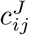 defines the coupling correlation between units *i* and *j*, and *δ*_*kk*_′ is the Kronecker delta function. In this setting, coupling correlation is imposed uniformly across all source units *k*, such that the similarity between *i* and *j*’s incoming couplings reflects a global alignment pattern, independent of the identity of the source unit. While this formulation abstracts away possible fine-grained heterogeneity in coupling correlations, it effectively constrains the structure of random neural networks with the similarity between incoming structural couplings across different units.

For standard DMFT to apply, an implicit assumption of weak coupling correlations must hold. In this section, we first work under this assumption and later examine its validity in biological data, as well as the consequences of its breakdown. Nevertheless, even weak coupling correlations prevent DMFT from being derived in exactly the same way as in conventional random networks with independent couplings. To “de-correlate” the system arising from structural couplings, we first introduce a latent coupling matrix **K** that satisfies **K** = **Λ**^−**1***/***2**^**U**^−**1**^**J**. Here, **U** = [**u**_**1**_,, · · · **u**_**N**_] and **Λ** = diag(*λ*_1_, · · ·, *λ*_*N*_) are the eigenvector and eigenvalue matrices of the coupling correlation matrix **C**, with entries 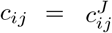, so that **C** = **UΛU**^−**1**^. After left-multiplying both sides of equation (1) by **U**^−**1**^, the dynamics become

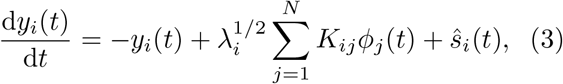

where *ϕ*_*j*_(*t*) abbreviates *ϕ*[*x*_*j*_(*t*)]. This linear transformation naturally introduces the latent dynamics **y**(**t**) = [*y*_1_(*t*), · · ·, *y*_*N*_ (*t*)] coupled through **K**, which are related to the original dynamics **x**(**t**) by **y**(**t**) = **U**^−**1**^**x**(**t**), with the external drive transformed as **ŝ**(**t**) = **U**^−**1**^**s**(**t**).

The entries *K*_*ij*_ remain Gaussian random variables with the same mean and variance as *J*_*ij*_, but they are mutually independent, satisfying E[*K*_*ij*_*K*_*mn*_] = *δ*_*im*_*δ*_*jn*_*g*^2^*/N* . Thus, by construction, coupling correlations are eliminated in the latent system.

More specifically, equation (3) describes an inhomogeneous system regarding *y*_*i*_(*t*) in which the factor 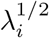 acts as a unit-specific scaling of the incoming coupling strength of latent unit *i*, introducing heterogeneity across the latent population. Nevertheless, apart from this coefficient 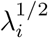, the form of equation (3) closely parallels that of equation (1) for conventional random neural networks with independent couplings and can therefore be analyzed using the same DMFT framework. As a result, the DMFT equation for the latent dynamics reads

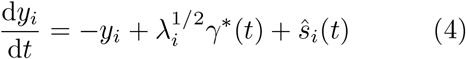

where *γ*^∗^(*t*) is a Gaussian stochastic mean-field term with self-consistent statistics ⟨*γ*^∗^(*t*)⟩_*t*_ =0 and ⟨*γ*^∗^(*t*)*γ*^∗^(*t*′)⟩_*t*_ = *g*^2^ ∑_*i*_⟨*ϕ*_*i*_(*t*)*ϕ*_*i*_(*t*′)⟩*/N* . Notably, the mean-field term *γ*^∗^(*t*) of different units is independent. Finally, transforming equation (4) back to the original coordinates via **x** = **Uy** yields the DMFT equation for the original system:

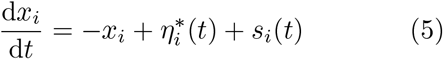

where 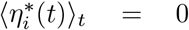 and 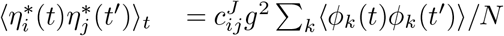.

To summarize, we first consider random neural networks with weak and non-zero coupling correlations, as defined in equation (2), and derive the corresponding DMFT equation (5), in which each unit is effectively governed by the Gaussian stochastic mean-field term 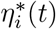, replacing the collective network interactions *η*_*i*_(*t*) in equation (1). The details of the DMFT equation derivation are provided in **Theoretical analysis of network dynamics** part of **Methods** and in **Supplementary information**.

## 3 DMFT predicts a linear relationship between coupling and dynamical correlation

From the DMFT equation, the dynamics *x*_*i*_(*t*) can be expressed as a stochastic integral driven by the mean-field fluctuation 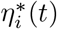 and the external input *s*_*i*_(*t*). To isolate the contribution of recurrent interactions that represent how coupling structure affects dynamics, we here set *s*_*i*_(*t*) = 0, so that the activity is entirely determined by internal coupling. In this case, *x*_*i*_(*t*) is a linear functional of the Gaussian input 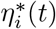, and its pairwise dynamical correlation depends solely on the cross-correlation structure of *η*^∗^. Defining the pairwise dynamical correlation as

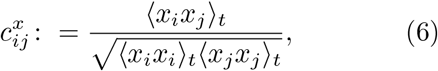

we obtain from equation (5) the exact identity

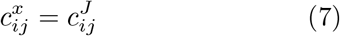

Thus, the emergent dynamical correlation 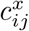 of network activity precisely mirrors the coupling correlation 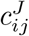 through a linear function with a unity slope. In other words, changes in dynamical correlation track changes in coupling correlation at exactly the same rate.

Moreover, according to equation (5), in the more biologically realistic setting where units receive external drive that may represent sensory inputs or subcortical signaling, the relationship between coupling and dynamical correlation is still linear, with the rate of change of dynamical correlation being suppressed and smaller than that of coupling correlation (**Methods**). Therefore, the DMFT theory establishes a rigorous linear mapping from coupling correlations to dynamical correlations, highlighting structural correlations as the key determinant of inter-unit synchrony.

It is worth noting that the DMFT prediction holds at the disorder-average level: the linear mapping between coupling and dynamical correlation is valid for the average across many unit pairs with the same coupling correlation, but not for any specific pair in a certain network realization. To test the theory, we therefore generate multiple random network realizations in numerical simulations and compute the average across them. Across different network realizations, the coupling correlation matrix **C** is held fixed, while the entries *J*_*ij*_ of the coupling matrix are generated as jointly Gaussian random variables with correlation specified by **C** (**Fig.2a**). To construct the coupling correlation matrices **C**, we use the sample correlation matrix of *N* Gaussian random variables (see **Numerical simulations** in **Methods**). This approach guarantees a symmetric, positive-definite, and statistically unbiased matrix, with correlation values randomly and weakly fluctuating around zero without any imposed structure.

**Fig. 2.**
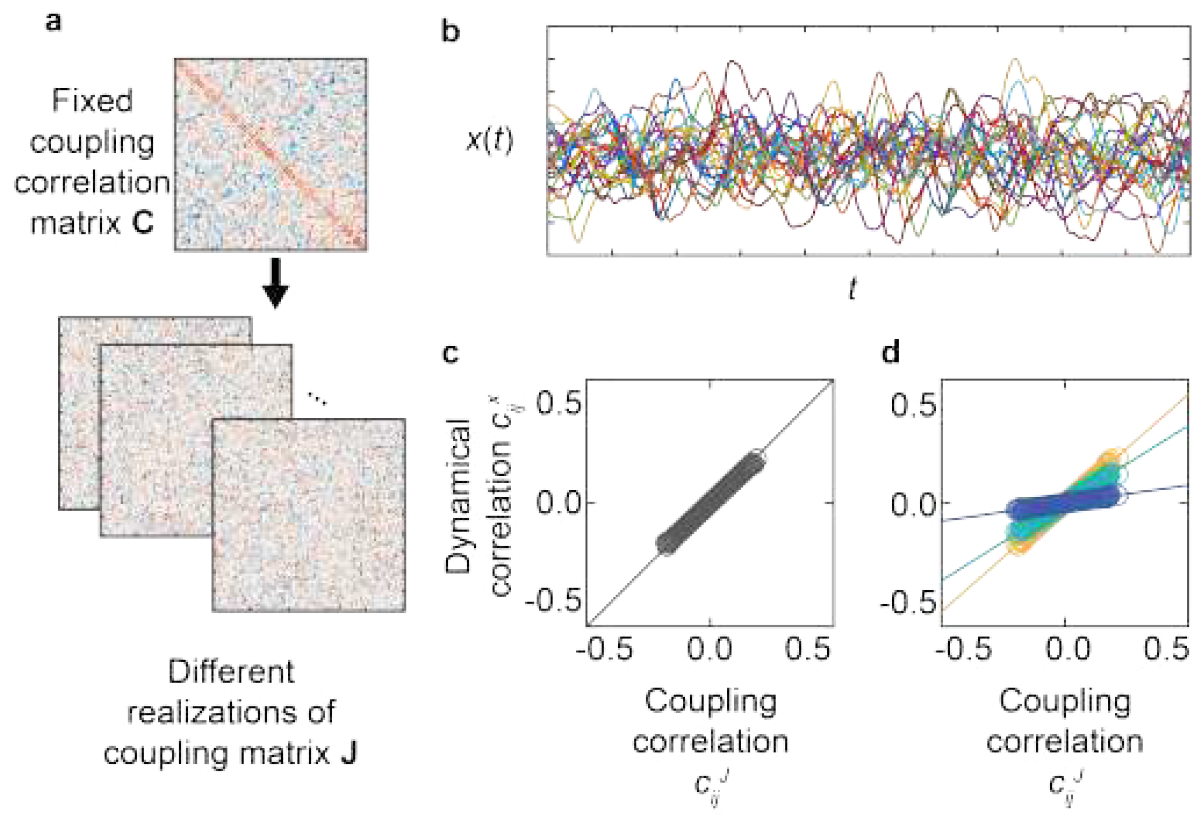
Numerical simulations of random neural networks with coupling correlations. **a**, Procedure for generating correlated random neural networks. A coupling correlation matrix **C** is first constructed and fixed. The coupling matrices **J** are then sampled from a multivariate Gaussian distribution whose covariance is determined by **C**. Consequently, different network realizations share the same coupling correlation while each possesses a unique coupling matrix. **b**, Example of chaotic dynamics exhibited by correlated random neural networks. **c**, In the absence of external drive, simulation results (open circles) precisely match the DMFT prediction 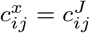 (solid line). **d**, Under non-zero external drive, the relationship remains linear, but the slope decreases with increasing drive magnitude.

The dynamics are simulated according to equation (1), with the external drive modeled as Gaussian white noise with a magnitude of ⟨*s*_*i*_(*t*)*s*_*i*_(*τ*)⟩_*t*_ = *δ*(*t* − *τ*)*σ*_*s*_. We ensure that networks operate in the chaotic regime (**Fig.2b**) by numerically estimating the maximal Lyapunov exponents (see **Methods**). This choice ensures that the dynamics remain non-stationary and exhibit sustained fluctuations, avoiding trivial fixed-point or decaying activity.

In the absence of an external drive, the simulation results (**Fig.2c**) show that any change in the coupling correlation (*x*-axis) produces an increase or decrease in the dynamical correlation (*y*-axis) with equal magnitude, precisely consistent with the theoretical prediction of 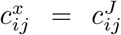. When nonzero *s*_*i*_(*t*) is introduced, the linear relationship is preserved, but its slope decreases, indicating that changes in dynamical correlations remain proportional to, yet smaller than coupling correlations (**Fig.2d**). These results align well with the theoretical predictions.

Therefore, the DMFT predicts a linear mapping between coupling and dynamical correlations, which is confirmed by numerical simulations using coupling correlation matrices constructed from the sample correlations of Gaussian random variables.

## 4 Biological coupling correlations exhibit long-tailed spectrum

However, the assumption of weak correlations and the above numerical simulations fail to capture important properties regarding the magnitude of the coupling correlation in the data. First, in simulations, the weak coupling correlation spans a narrow range, with absolute values typically below 0.1 ∼ 0.2 (*N* = 500, **Fig.3a**), while in biological networks, the magnitude of the coupling correlation could approach or exceed 0.5 (**Fig.3c-e**). Furthermore, in numerical simulations, the coupling correlation magnitude diminishes further as the system size increases (**Fig.3b**). By contrast, biological networks show a large coupling correlation magnitude that is roughly invariant across species and levels of network granularity (**Fig.3b**, inset).

**Fig. 3.**
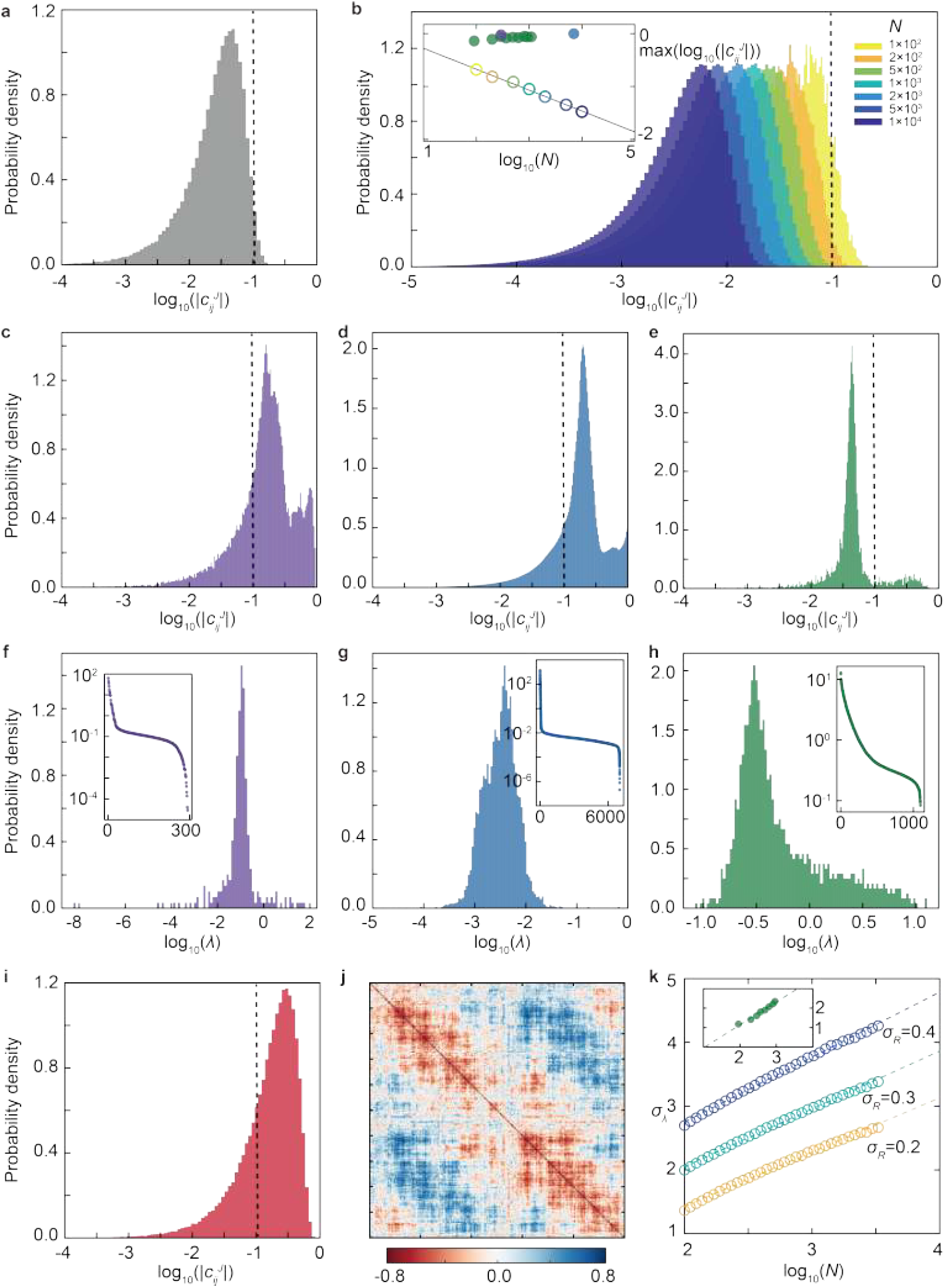
The spectrum and the coupling correlation magnitude. **a**, Distribution of the absolute coupling correlation values in simulated networks with bulk spectra. The *x*-axis represents the logarithm of absolute 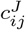, and the vertical dashed ne marks a correlation magnitude of 0.1. **b**, As system size *N* increases, the distribution of coupling correlation magnitudes shifts leftward, indicating a decrease in overall correlation strength. The inset shows that the maximum coupling correlation scales inversely with *N* ^1*/*2^, appearing as a line with slope − 1*/*2 on the log–log plot (empty circles). In contrast, biological networks exhibit size-invariant maximum coupling correlation magnitudes across species (*Drosophila* in purple, mouse in blue, human in green). **c–e**, Distributions of coupling correlation magnitudes for *Drosophila*, mouse, and human data, respectively. **f–h**, Spectra of the coupling correlation matrices for the same datasets, where the *x*-axis denotes the logarithm of eigenvalues *λ*. The spectra span several orders of magnitude and appear bulk-like only on a logarithmic scale. Insets show the sorted spectra, with *y*-axis representing eigenvalues and *x*-axis their rank. **i**, Distribution of coupling correlation magnitudes in simulations with long-tailed (log-normal) spectra, which successfully exhibit coupling correlations with large magnitude. **j**, Example of a simulated coupling correlation matrix, with rows and columns sorted to reveal patterned structure. **k**, Scaling of the log-normal spectral parameter *σ*_*λ*_ (*y*-axis) with system size *N* (*x*-axis), which is required for the correlation magnitude to remain invariant. Specifically, *σ*_*λ*_ scales weakly with *N*, manifesting as a proportionality o log_10_(*N*). Empty circles denote simulation results for different fixed correlation magnitudes (defined by the standard deviation of the coupling correlation matrix). The inset shows the corresponding human data, where *σ*_*λ*_ increases linearly with log_10_(*N*), consistent with the simulation results.

This discrepancy arises from the eigenspectrum properties of the coupling correlation matrix. In the previous simulated networks, the spectrum of the coupling correlation matrix is characterized by a bulk distribution in which all eigenvalues lie within a compact range and are of comparable orders of magnitude (**Methods** and **Supplementary Fig.1**). Random matrix theory predicts that in such bulk spectrum, the scale of the correlation matrix decays as *N* ^−1*/*2^, consistent with the vanishing correlation magnitude in simulations (**Fig.3b**). In biological networks, however, finite and size-invariant correlation magnitude implicates a long-tailed spectrum with eigenvalues spanning multiple magnitudes – many small, near zero, and few very large. Indeed, biological coupling correlation matrices exhibit such long-tailed spectra, appearing bulk-like only logarithmically (**Fig.3f-h**). In fact, numerically generated coupling correlation matrices with randomly sampled long-tailed spectra successfully exhibit biologically plausible correlation magnitudes (**Fig.3i**). Interestingly, such coupling correlation matrices are highly structured (**Fig.3j**), unlike the case of bulk spectra, where the correlation matrix appears noisy (**Fig.2a**).

We then quantitatively investigate the condition for coupling correlation to be size-invariant using simulation (**Methods** and **Supplementary Fig.2**). Specifically, the eigenvalues are drawn from log-normal distributions—consistent with the empirical spectra observed in biological data—characterized by a single free parameter, *σ*_*λ*_. The other parameter of the log-normal distribution, *µ*_*λ*_, is not treated as independent but is constrained by 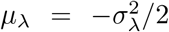, ensuring that the eigenvalues satisfy the normalization condition ∑_*i*_ *λ*_*i*_*/N* = 1, as required for correlation matrices. Here, the parameter *σ*_*λ*_ serves as the dispersion parameter of the log-normal spectrum, governing the spread and tail heaviness of eigen-values in logarithmic space. Larger *σ*_*λ*_ produces a more heterogeneous spectrum with a heavier tail, corresponding to stronger coupling strength heterogeneity in the latent population. The typical magnitude of the generated correlation matrices is quantified by the standard deviation of their entries, denoted by *σ*_*C*_. Simulation results show that for *σ*_*C*_ to remain fixed, *σ*_*λ*_ must scale weakly with system size *N*, being proportional to its logarithm (**Fig.3k**). This result closely matches the experimental data (**Fig.3k**, inset). Therefore, a biologically plausible coupling correlation matrix—exhibiting an appropriate magnitude and remaining invariant to system size—is realized by a log-normal spectrum whose dispersion parameter *σ*_*λ*_ scales only weakly with *N*, as revealed by both simulations and empirical data.

## 5 Long-tailed spectrum yields an approximately linear relationship between structural and functional correlation

The spectral property not only explains the discrepancy in coupling correlation magnitudes, but also has non-trivial implications for network dynamics. While in previous simulations, the networks with weak coupling correlations are characterized by a bulk spectrum; here, we focus on the dynamics in networks that exhibit long-tailed spectra. The spectra are sampled from log-normal distributions, with *σ*_*λ*_ determined by the network size *N* and the desired coupling correlation magnitude *σ*_*C*_. In the absence of an external drive, the results reveal a moderate deviation from linearity: across most values of coupling correlation, the relationship between coupling and dynamical correlation remains nearly linear, whereas noticeable nonlinearities arise only at strong coupling correlations (**Fig.4a**). Under the external drive, similar deviations occur—the relationship departs from the theoretical linear function predicted for bulk spectra, exhibiting approximate linearity with increasing nonlinearity as the coupling correlation magnitude approaches unity (**Fig.4b-d**).

**Fig. 4.**
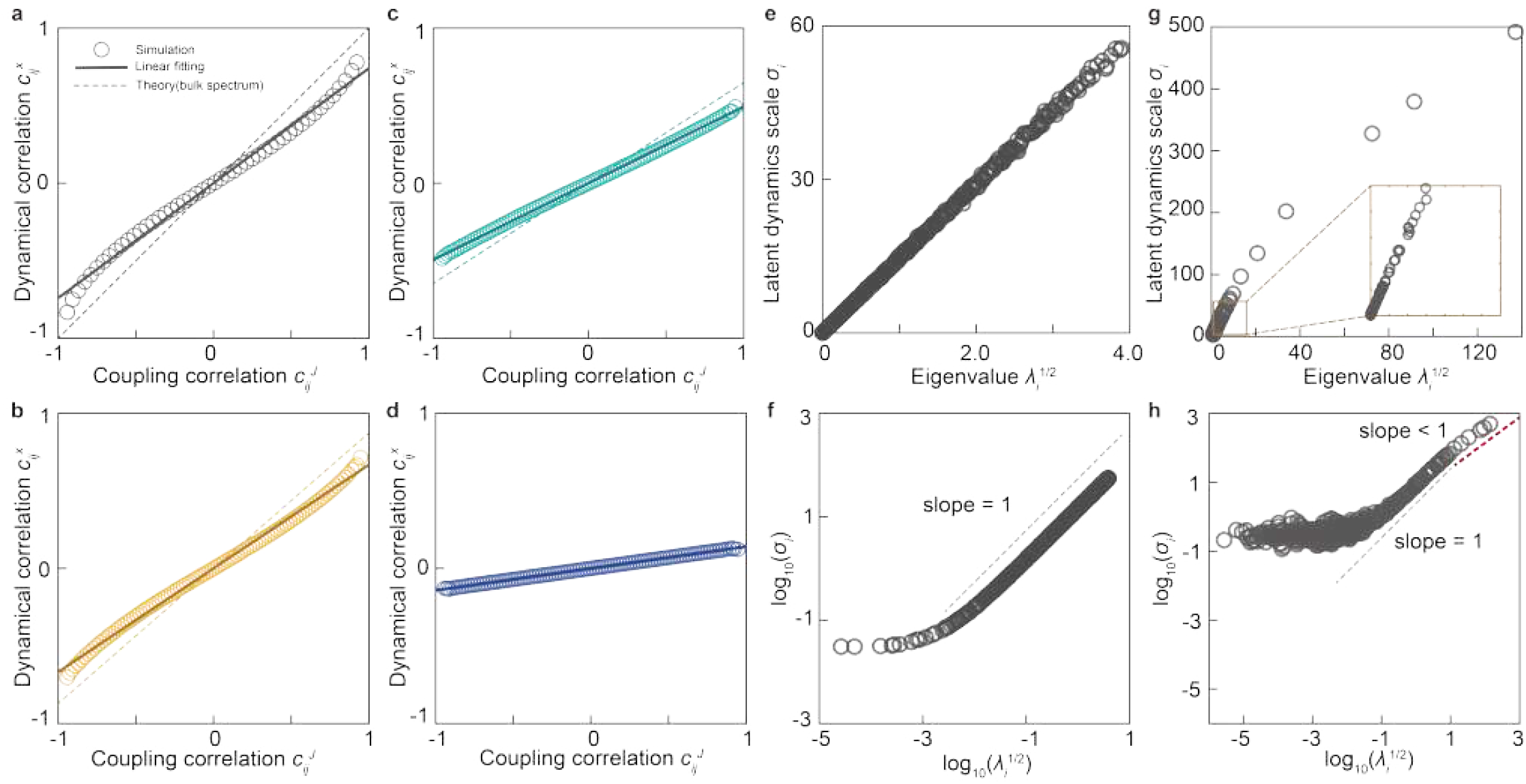
Long-tailed spectra influence network dynamics. **a**, Simulated relationship between coupling correlation (*x*-axis) and dynamical correlation (*y*-axis) in the absence of external drive under long-tailed spectra. Empty circles denote simulation results, the solid line indicates the best linear fit, and the dashed line shows the DMFT prediction for bulk spectra case. The scatter can be well approximated by a linear trend with a smaller slope than the DMFT prediction, while noticeable nonlinearities emerge only when coupling correlation approaches unity. **b–d**, Results for non-zero external drive modeled as Gaussian white noise with standard deviation 10^1^, 10^1.5^, and 10^2^, respectively. **e**, For bulk spectra, the scale *σ*_*i*_ (*y*-axis) of simulated latent dynamics *y*_*i*_(*t*) is proportional to the corresponding eigenvalue 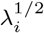 (*x*-axis). **f**, The same results as in **e** but shown on a log–log scale, where the linear relationship between *λ*_*i*_ and *σ*_*i*_ is evident from the unity-slope line. **g**,**h**, Relationship between 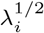 and *σ*_*i*_ for long-tailed spectra, plotted in the same way as in **e**,**f**. In **g**, the relationship appears linear for small *λ*_*i*_ (see inset) and nonlinear for large *λ*_*i*_. A clearer view of the complex nonlinear relationship is provided by panel **h**: a plateau for very small *λ*_*i*_, a linear regime indicated by unity slope on log-log plot (gray dashed line) for intermediate *λ*_*i*_, and a sublinear regime for large *λ*_*i*_ indicated by a slope smaller than one (red dashed line).

These deviations reflect the implicit assumption of weak correlations in the DMFT derivation. Specifically, we drew an analogy between the latent dynamics of correlated random neural networks and the dynamics of conventional fully random networks. The validity of this analogy critically relies on the pre-scaling term *γ*_*i*_(*t*) = ∑_*j*_ *K*_*ij*_*ϕ*_*j*_(*t*) in equation (3) being homogeneous in the thermodynamic limit, as is the case for *η*_*i*_(*t*) in conventional random neural networks. In other words, for different units *i*, their incoming collective interactions, represented by *γ*_*i*_(*t*), must converge to a common mean-field term *γ*^∗^(*t*). This homogeneity justifies the saddle-point approximation in the DMFT derivation (**Methods** and **Supplementary Information**), which assumes the existence of a single, well-defined “most probable” state for the collective interaction *γ*_*i*_(*t*). Such an assumption holds in networks with bulk spectra, where correlations between *ϕ*_*i*_(*t*) are weak. However, for long-tailed spectra, strong coupling correlations induce strong dependencies among *ϕ*_*i*_(*t*), violating the mean-field convergence of *γ*_*i*_(*t*). In this regime, *γ*_*i*_(*t*) may remain highly heterogeneous, with its statistics varying from unit to unit, depending on their position within the network structure. Consequently, the system’s macroscopic state is no longer described by a single saddle point but rather by a distribution of effective fields, leading to the failure of the standard DMFT prediction.

This picture of heterogeneity-driven failure is directly supported by our numerical analysis of the latent dynamics. To assess the validity of the homogeneity condition under different spectral distributions, we numerically examined the relationship between the scale of latent dynamics *y*_*i*_(*t*), quantified by its temporal standard deviation *σ*_*i*_, and the corresponding eigenvalue *λ*_*i*_. According to equation (3), *σ*_*i*_ should change linearly with the scale of 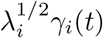. Therefore, a proportional relationship between *σ*_*i*_ and 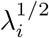 ports the DMFT assumption, whereas systematic deviations from this linearity indicate a failure of the mean-field approximation due to persistent heterogeneity in *γ*_*i*_(*t*).

After simulating the original network dynamics **x**(**t**) with equation (1), we compute the latent dynamics **y**(*t*) as **y**(*t*) = **U**^−**1**^**x**(*t*). When *λ*_*i*_ follows a uniform distribution—corresponding to a bulk spectrum—the latent dynamics scale *σ*_*i*_ is indeed proportional to 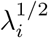, as indicated by the straight line of unit slope on the log–log plot (**Fig.4e,f**). In contrast, when *λ*_*i*_ is drawn from a long-tailed distribution, the relationship between *σ*_*i*_ and 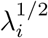 deviates markedly from linearity: as shown in **Fig.4g,h**, the scale of latent dynamics saturates for small *λ*_*i*_, followed by linear scaling with 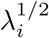. Finally, it grows sublinearly with 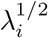 for large *λ*_*i*_, corresponding to a slope smaller than one on the log–log scale.

This correspondence between the linearity (or nonlinearity) of the 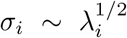 relationship and that of the 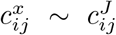 relationship persists across different spectral distributions: linearity of both relationships holds as long as the spectrum is bulk-like, and breaks down otherwise (**Supplementary Fig.3&4**). Therefore, the simulations demonstrate that when the coupling correlation exhibits a long-tailed spectrum, the collective interaction term *γ*_*i*_(*t*) fails to converge to a common mean-field term *γ*^∗^(*t*), thereby explaining the observed deviation from the DMFT-predicted structure–function relationship.

Nevertheless, the deviation from DMFT prediction is not fundamental. Specifically, in **Fig.4a-d**, when the absolute coupling correlation approaches unity, the corresponding dynamical correlation increases superlinearly; conversely, when coupling correlations are not particularly large, the relationship remains approximately linear but with a slope smaller than one, indicating that dynamical correlations vary more slowly with coupling correlations than previously predicted. Meanwhile, the deviation from linearity is further suppressed by external inputs (**Fig.4c-d**). This is consistent with biological data, where coupling correlations rarely reach extreme values, external inputs exist, and the relationship between coupling and dynamical correlations is well captured by a linear approximation **Fig.6a-c**, as will be shown in the next section. More importantly, the approximate linearity ensures that the scale of dynamical correlation matches that of coupling correlation, so that long-tailed spectra confer scale invariance not only to structural coupling correlations but also to the emergent dynamical correlations in large systems. This robust correlation scale enables the coordinated activation of different units—a feature that is likely critical for supporting coherent brain function in large, biological networks.

## 6 Validation with biological data

The numerical simulations in the previous section revealed that deviations from linearity are confined to coupling correlations approaching unity in magnitude, and are further suppressed in the presence of external inputs. This observation motivates a concrete prediction for brain data: if region pairs with extreme coupling correlations constitute only a small fraction of all pairs, and brain regions receive substantial external inputs beyond their interactions, then the structure-function relationship in biological neural networks should fall within the approximately linear regime, despite the formal breakdown of the DMFT assumption. Indeed, across *Drosophila*, mouse, and human datasets, only a small fraction of region pairs exhibit coupling correlations approaching unity, and brain regions are continuously subjected to external inputs from sensory systems and subcortical structures. These two conditions together place biological networks within the range where approximate linearity is expected to hold. The binning analysis below tests this prediction directly.

We analyze publicly available datasets from human cortex, mouse cortex, and *Drosophila* hemibrain, each containing both neural dynamics recordings (fMRI for human [26, 27]; wide-field calcium imaging for mouse [28] and *Drosophila* [29, 30]) and macroscopic anatomical connectivity information (DTI for human [26, 27]; bulk tracing for mouse [31, 32]; electron microscopy reconstruction for *Drosophila* [29, 33]). Consistent with our theoretical definitions, the dynamical correlation matrix is computed from the neural activity of brain regions as Pearson’s correlation coefficients (**Fig.5a**), while the coupling correlation matrix is computed from the anatomical connectivity matrix as the correlation coefficients between its rows (**Fig.5b,c**).

**Fig. 5.**
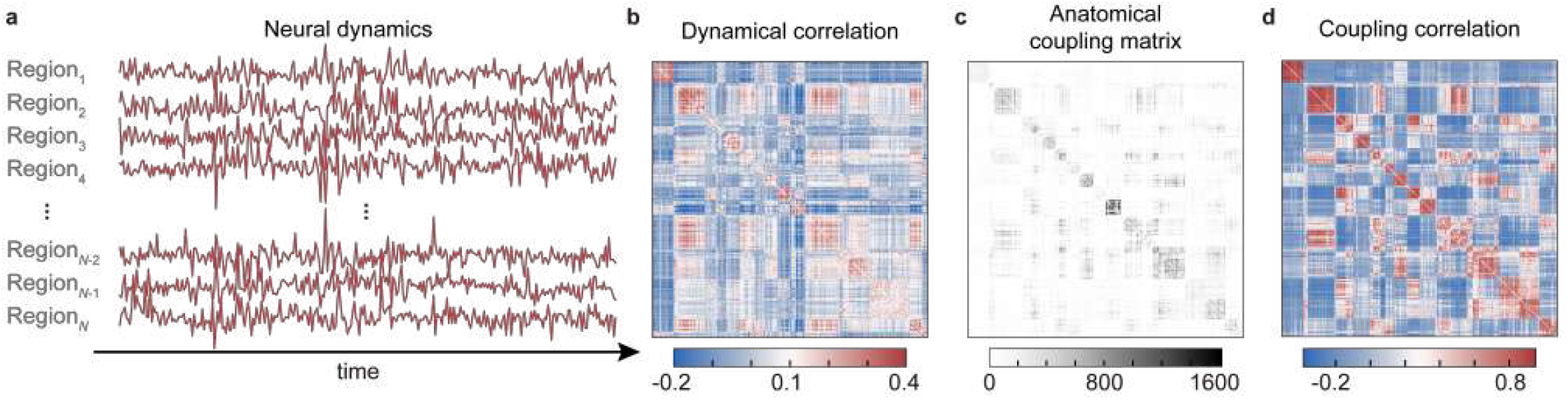
Coupling and dynamical correlation in biological data. **a**, Illustration of neural activity with *Drosophila d*ata. **b**, Illustration of dynamical correlation matrix with *Drosophila*. Each element in the dynamical correlation matrix represents the Pearson correlation coefficient between the neural activity of regions *i* and *j*. **c**, Illustration of anatomical coupling matrix with *Drosophila* data. **d**, Illustration of coupling correlation matrix with *Drosophila* data. Each element the coupling correlation matrix is computed as the Pearson correlation coefficient between the *i*-th and *j*-th rows of the anatomical coupling matrix.

The DMFT prediction applies to the disorder average. However, since we have only a single network realization (i.e., one brain) for each species, it is impractical to average over multiple realizations as in the numerical simulations. To overcome this, we leverage the large number of node pairs within a single network, performing a binning analysis with fixed intervals in coupling correlation. Specifically, we divide the experimental range of coupling correlation into *N*_bin_ bins with equal width. Within each bin, we compute the average values of coupling and dynamical correlation, which provides an empirical estimate of the expectation of dynamical correlation given coupling correlation.

This approximation relies on a self-averaging assumption: in large and heterogeneous networks, averaging over node pairs within a single realization provides a proxy for the disorder average. Here, heterogeneity refers to the broad distribution of coupling correlations across node pairs, which is empirically observed in all datasets (**Fig.3c-e**). Also, the resulting binning analysis is stable across binning granularities (see below in **Fig.6j-l**), supporting the validity of this approximation.

**Fig. 6.**
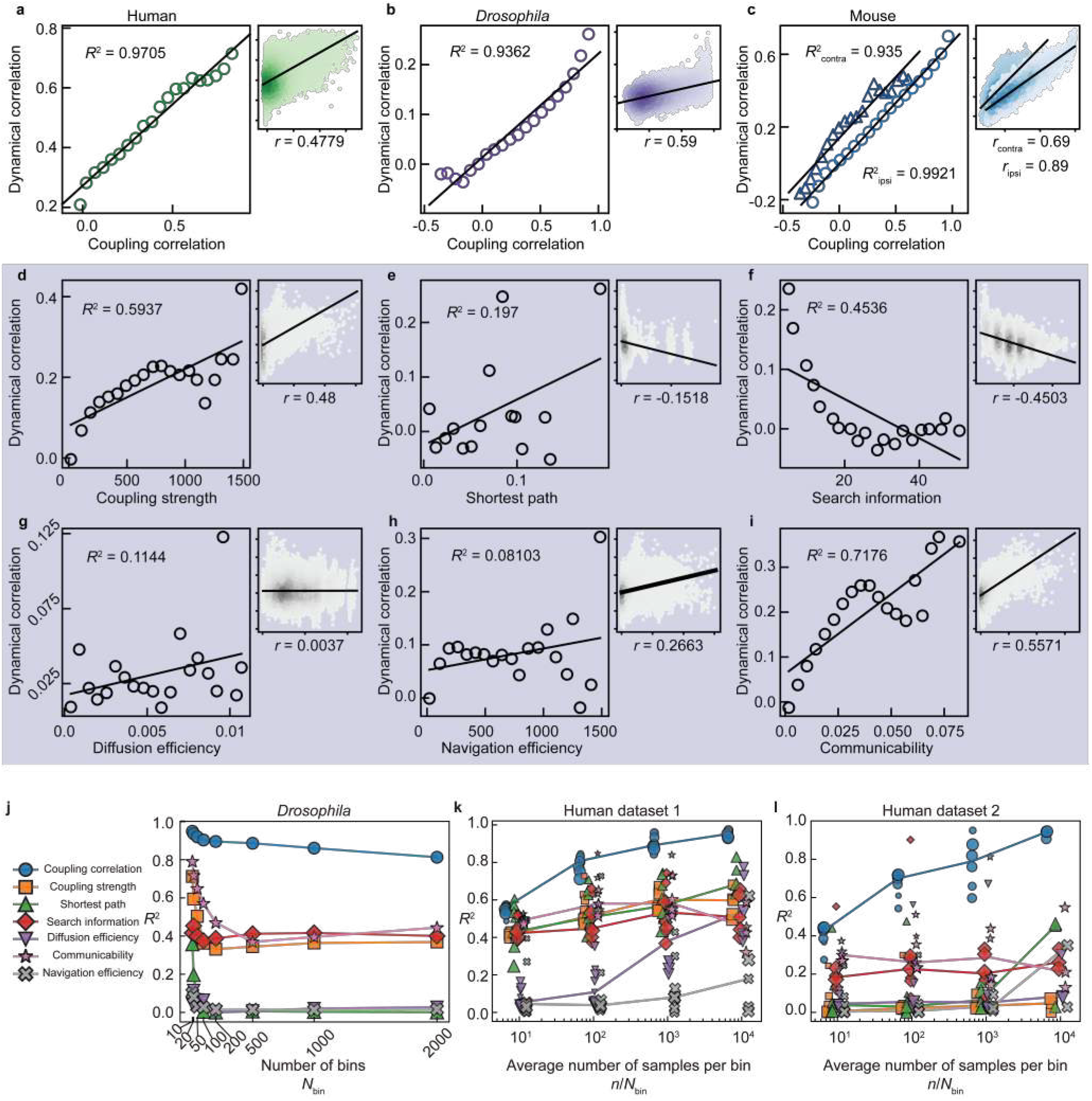
Coupling and dynamical correlation exhibit a linear relationship. **a**, Scatter plots of dynamical correlation (*y*-axis) versus coupling correlation (*x*-axis) in human data. the main (left) panel shows binned relationship between coupling and dynamical correlation (circles) with *N*_bin_ = 20. The side (right) panel shows the raw scatter plot, with color intensity indicating data-point density (darker colors denote higher density). The black solid lines indicate linear fits. *R*^2^ represents the coefficient of determination of linear fitting, and *r* is the Pearson correlation coefficient. **b**, Same as **a** for *Drosophila* data. **c**, Same as **a** for mouse data, analyzed separately for contralateral (darker blue, triangles) and ipsilateral (lighter blue, circles) region pairs. Note that in the sider panel if the contralateral and ipsilateral data are not visualized separately, their superposition gives rise to a bump of scatters that deviates from a overall linear trend. **d-i**, The results of direct coupling strength (**d**) and five network communication measures (shortest path, search information, diffusion efficiency, navigation efficiency, communicability) of *Drosophila* data at *N*_bin_ = 20. **j**, The linear fitting *R*^2^ of between binned dynamical correlation and different structural features of *Drosophila* data under a wide range of *N*_bin_. **(k)** Linear fitting *R*^2^ between binned dynamical correlation and various structural features for human dataset 1 (HCP data). Markers of identical shape but varying size correspond to different brain parcellation granularities: larger markers indicate parcellations with more regions. For each parcellation, results are obtained with a different number of bins *N*_bin_, chosen such that the average number of samples per bin *n*_avg_ remains the same across all parcellations. Lines show how the median *R*^2^ across parcellations changes with *n*_avg_. **(l)** Same as **(k)** for human dataset 2 (MICA data). Here, markers of increasing size denote five parcellation schemes (ordered from smallest to largest marker): Van Economo cytoarchitectonic atlas, Destrieux sulco-gyral atlas, Glasser multimodal atlas, fine-grained subparcellations of the Desikan-Killiany atlas, and three Schaefer functional atlases with varying node counts. Multiple Schaefer resolutions are included to provide multiple samples for the case where *n*_avg_ = 10^4^. Other parcellation schemes cannot be used for this case, as they do not provide a sufficient number of brain regions.

The binning analysis of human, *Drosophila* and mouse data reveals results consistent with the theoretical predictions (**Fig.6a-c**). Before binning, coupling and dynamical correlations are positively correlated across all three species, and the scatter plots exhibit a discernible linear trend (side panels of **Fig.6a-c**). After binning, the relationship becomes strongly linear, with a coefficient of determination *R*^2^ *>* 0.9 for each species (main panels of **Fig.6a-c**, *N*_bin_ = 20). Interestingly, the raw scatter plot of the pre-binning mouse data—which include coupling and dynamical correlations from both brain hemispheres—displays an anomalous pattern where a localized “bump” of points appears to deviate from the overall linear trend. However, when the ipsilateral and contralateral data are visualized and analyzed separately, this anomaly resolves cleanly: each subset on its own exhibits a clear linear trend before binning and a strongly linear relationship after binning.

We further compared coupling correlation with direct coupling strength and five popular network communication measures [17]: shortest path, search information, diffusion efficiency, communicability and navigation efficiency, first using *Drosophila* data for test (**Fig.6d-i**). In the side panels, several of these measures show a statistically significant correlation with dynamical correlation. However, their correlation coefficients are consistently lower than that of coupling correlation. More importantly, we performed a binning analysis to test whether these statistical correlations reflect a genuine linear causal relationship at the disorder-average level (main panels of **Fig.6d-i**, *N*_bin_ = 20). Some measures exhibit no linear relationship, with binned points scattered randomly and an *R*^2^ near zero. Others yield a higher *R*^2^ but still substantially lower than coupling correlation, and display clear nonlinear (e.g., search information), even non-monotonic (e.g., communicability), twists in their binned relationship—reminiscent of the ad hoc nonlinear transformations (e.g., logarithmic or inverse scaling) employed in previous studies. These results indicate that the relationship between these measures and dynamical correlation is unlikely to be both causal and linear at the disorder-average level. They are either causal but fundamentally nonlinear (requiring arbitrary transformations), or linear but merely correlational (with twists reflecting statistical noise). In either case, therefore, linear regressions based on direct coupling strength or standard communication measures do not provide a direct mechanistic link. In stark contrast, our coupling correlation reveals a clean, theoretically grounded linear relationship at the disorder-average level, empirically verified by the data.

The *Drosophila* results are robust across binning granularities: when *N*_bin_ ranges from 10^1^ to 10^3^, coupling correlation consistently exhibits a strong linear relationship with dynamical correlation, with an *R*^2^ that remains higher than that of all other measures (**Fig.6j**). We did not perform this comparison on mouse data because the coupling matrix contains thousands of nodes which makes the computation of network communication measures prohibitively time-consuming. Instead, we examined human data using two independent datasets. The first dataset, derived from Human Connectome Project (HCP) data [26], provides brain parcellations with region counts ranging from 10^2^ to 10^3^ using a consistent parcellation method (Schaefer atlas). The second dataset, based on Multimodal Imaging and Connectome Analysis (MICA) data [27], uses multiple parcellation methods, each yielding a different number of brain regions. The accuracy of the disorder-average approximation depends on the number of samples averaged per bin. We therefore define *n*_avg_ = *n/N*_bin_ as the average samples per bin, and vary it from 10^1^ to 10^4^. For each parcellation (with total region pairs *n*), the number of bins *N*_bin_ is then determined accordingly. The linear fitting *R*^2^ for the first human dataset is shown in **Fig.6k**. Markers of different sizes correspond to different parcellation resolutions, and the lines represent the median *R*^2^ values across resolutions. The *R*^2^ for coupling correlation (blue circles) is consistently higher than that of direct coupling strength and all communication measures, approaching 0.9 − 1.0 when *n*_avg_ reaches the order of 10^2^ to 10^3^. A similar phenomenon is observed in the second human dataset (**Fig.6l**), which provide the results of brain parcellation atlases guided by distinct principles, for example, cytoarchitecture, sulco-gyral landmarks, functional and mutimodal brain imaging data.

Therefore, the experimental data support the first-principles DMFT prediction that coupling correlation exhibits a clean linear relationship with dynamical correlation at the disorder-average level. In contrast, although prevalent network communication measures may capture certain aspects of network topology and show statistical correlation with dynamical correlation, their predictive performance for dynamical correlation is quantitatively inferior to that of coupling correlation. Moreover, they lack a causally grounded, analytically derived link between structure and dynamics—a gap that our framework aims to fill.

To summary, our study provides a mechanistic and quantitative account of how brain structure dictates dynamics across scales. First, combining theoretical analysis with DMFT and numerical simulation on random neural network models, we identify coupling correlation as the structural determinant of dynamical correlation. Second, our analysis reveals that their relationship, while dependent on spectral properties, is predominantly linear—precisely so for bulk spectra and approximately for the long-tailed spectra that confer biological realism. Finally, this predicted approximate linearity is robustly confirmed by diverse biological data.

## 7 Discussion

Our findings reveal that the coupling correlation provides an analytically tractable structural predictor of inter-unit dynamical correlations in large-scale networks. Behind this relationship lies a simple yet powerful principle: behavioral similarity between two units arises from the similarity in their received inputs, which in turn reflects their structural similarity. In large-scale neural networks, this structural similarity is captured by the correlation between rows of the coupling matrix—quantifying how similar the input connectivity patterns of different units are.

Supporting evidence for this principle appears across biological systems of different scales. In C.*elegans*, for example, dynamical correlations between single-neuron activities are shaped by the fraction of shared neighbors between neurons—a non-local structural feature analogous to coupling correlation in our framework, though defined in a discrete setting [34, 35]. Similarly, in the transcriptional regulatory network of E.*coli*, genes that share isomorphic input trees—the set of regulatory pathways terminating at each gene—exhibit synchronized expression patterns that sustain cellular functions [36]. There is, in fact, mounting evidence that both synaptic and gene-regulatory networks exhibit systematic deviations from randomness, as revealed by motif analyses. In binarized or otherwise discretized representations of these networks, divergent motifs—triplets in which a single unit sends outputs to two others—are consistently over-represented [37–43]. The prevalence of such motifs potentially provides a unifying structural description across systems: in discrete networks, they reflect shared input or output patterns, while in weighted or continuous networks they may manifest as correlated structural couplings, as in our formulation. Together, these findings suggest that non-random, motif-based structure—and its continuous analogue, coupling correlation—is a ubiquitous organizational principle of biological networks that shapes their emergent dynamics.

Here, our work provides probable theoretical framework that is lacking in previous studies: we not only provide a measure of structural and dynamical similarity with coupling and dynamical correlation, but also establish their quantitative relationship that is approximately linear. Beyond neural systems, by replacing random neural networks with other suitable models, the concept of coupling correlation and the resulting structure–function mapping may be extended to a broad class of complex networked systems.

The magnitude of coupling correlation and its size-invariance is not a trivial observation and is closely related to the spectral property. Wang et al. [44] found a brain-wide scale-invariant spectrum in the covariance matrix of neural dynamics, characterized by a long-tail eigenvalue distribution. Our analysis identified a dual long-tailed spectrum via structural correlations. Coherent neural activity has been modeled by embedding low-rank connectivity in complex networks [45–54]. Unlike typical low-rank systems with a few dominant eigenmodes, our long-tailed spectrum maintains a broad hierarchy of eigenvalues, leading to highly correlated networks. This structure may offer functional benefits similar to low-rank organization, such as efficient control and resilience, and may explain the prevalence of structured organization in natural and artificial networks, suggesting a fruitful area for further study.

Finally, several open questions remain for future work. First, in this study, for coupling correlations with long-tailed spectra, the dynamical properties—including chaos and dynamical correlations—were examined purely through numerical simulations. While chaos can indeed be confirmed numerically, developing an analytical framework would offer deeper insight into the dynamics, uncovering distinct dynamical regimes and possible phase transitions as functions of network parameters. Moreover, there is also a need for a theoretical framework to describe the collective interaction term *γ*_*i*_(*t*) when it does not converge to a single mean-field term *γ*^∗^(*t*) in networks with long-tailed spectra, which could provide deeper insight into the nature of correlated network systems. Second, in the theoretical analysis of dynamics, we considered the case of a single steady state in which dynamical correlations are time-translation invariant. However, empirical studies have shown that animal brains exhibit distinct correlation patterns across behavioral states and tasks, whereas the assumption of single steady state results in a unique correlation pattern as long as the coupling and external drive remain unchanged. A plausible explanation is that inputs from subcortical nuclei or physiological signals vary over time, leading to shifts in dynamical correlation patterns. Further investigation is required to quantitatively characterize such external drives, and to explore whether random neural networks is possible to intrinsically transition between multiple steady states. Lastly, although the coupling strengths in our theory and simulations follow Gaussian distributions, biological neural networks typically display long-tailed coupling strengths [33, 55, 56]. Theoretical frameworks for analyzing long-tailed systems remain scarce [57, 58]. Moreover, defining coupling correlation becomes mathematically challenging, as long-tailed variables often exhibit divergent means or variances. Addressing these challenges will not only deepen our understanding of brain dynamics but also shed light on the behavior of complex network systems more broadly.

## 8 Acknowledgement

The study was supported by grants from National Science and Technology Major Program (Grant No. 2025ZD0219300 and No. 2022ZD0205000 to C.L.), the National Natural Science Foundation of China (32171088 and 32427802 to C.L.).

## 9 Author information

### 9.2 Contributions

C.L. and Qihang Wu identified the problem. Qihang Wu proposed the hypothesis, implemented the theoretical analysis, performed numerical simulations, analyzed the experimental data, and wrote the manuscript. C.L. and Quan Wen revised the manuscript and supervised the study.

## 11 Methods

### 11.1 Theoretical analysis of network dynamics

We follow the well-established theoretical framework of DMFT for analyzing the dynamical properties of Gaussian random neural networks, which requires three steps: 1) calculating the average generating functional of the dynamical process with the path integral method, 2) deriving the DMFT equations from the average generating functional via saddle-point approximation, and 3) analyzing the dynamical properties from the DMFT equations. In what follows we would mention some important intermediate results that promote understanding of the theoretical derivation. Other details of each step will be presented in the **Supplementary information**.

To construct the generating functional, we add a perturbation field *j*_*i*_(*t*) to the original system described by equation (1):

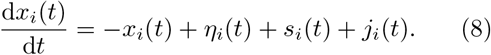

In the first step, we begin by calculating the generating functional for a specific network realization **J**, which is straightforward according to the definition:

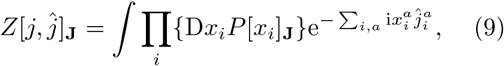

where 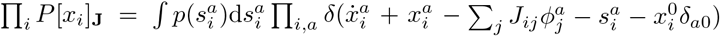 is the probability density of the dynamical path constrained by the differential equation (1) and the initial condition 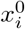, and 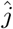 is a source field used for defining the generating functional. *x*^*a*^ represents *x*(*t*_*a*_) and ∑_*a*_ *x*^*a*^ abbreviates 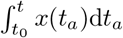. The conjugate variable 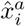 is introduced by applying the Fourier representation of Dirac *δ* function

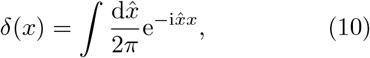

which finally results in the generating functional that reads:

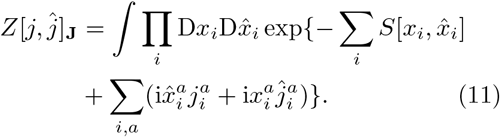

Here, 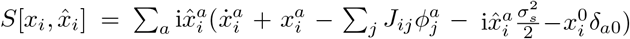 is called the action of the dynamical path and governs the dynamics through the path integral, where each dynamical path is weighted by 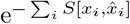. The path that minimizes *S* dominates the generating functional, while fluctuations around it are suppressed exponentially with increasing *S*.

Crucially, since the generating functional *Z*_**J**_ is dependent on the specific coupling matrix **J**, we then focus on the disorder-averaged (over all random coupling matrices) generating functional 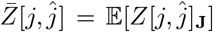 to extract universal features of dynamical correlations, determined solely by the statistics of **J** (e.g., coupling alignment) while being independent of specific coupling matrix realizations. This approach bridges microscopic disorder (e.g., coupling matrix realizations) and macroscopic observables (e.g., dynamical correlations), akin to studies of random neural networks and spin glasses. Given the multivariate Gaussian distribution of *J*_*ij*_, the average of *Z*[*j, ĵ*]_**J**_ over disorder gives

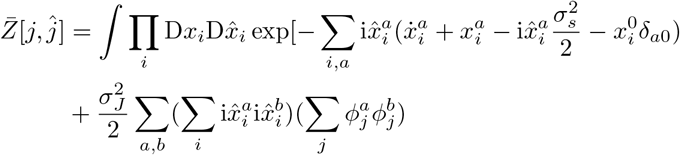

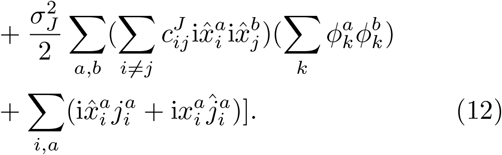

Notably, the presence of coupling correlation 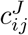 gives rise to the non-local interaction term 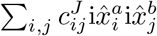, in contrast to conventional random neural networks where the statistical independence of *J*_*ij*_ causes the non-local interaction to vanish. Unfortunately, the construction of DMFT equations in the conventional cases is based on the approximating 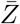 as the product of *N* generating functionals of respective stochastic processes, which strictly requires the absence of non-local interaction. Therefore, the reduction of this non-local interaction term is necessary.

As mentioned in the main results, this was achieved via the introduction of latent dynamics described by equation (3). By applying the analysis so far about equation (1) on equation (3), we give the generating functional for the latent dynamics that reads

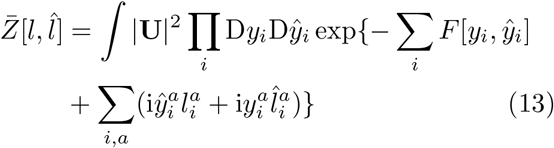

where 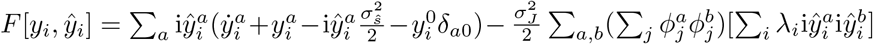 is the action of the latent dynamics, and |**U**| ^2^ = 1 is the determinant of **U**.

In the second step of deriving the DMFT equations, the saddle-point approximation of 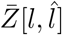 in the thermodynamic limit reads:

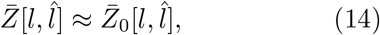

where 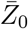 can be subsequently viewed as the product of *N* individual generating functionals of single-site stochastic dynamics:

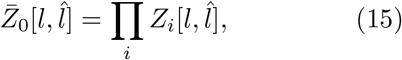

with

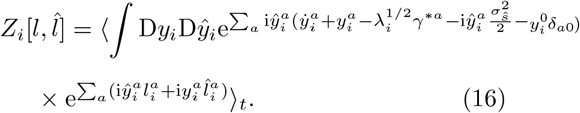

The derivation of the saddle-point approximation 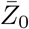 involves multiple steps and is presented in the **Supplementary information**.

Importantly, 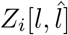 is the generating functional of the latent dynamics *y*_*i*_, which is in fact a stochastic process driven by the mean-field fluctuation 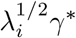. Finally, in the third step, the latent dynamics *y*_*i*_ can be described by the stochastic differential equation (4), which then gives the DMFT equation for *x*_*i*_ as equation (5). From equation (5) the solution of *x*_*i*_ reads

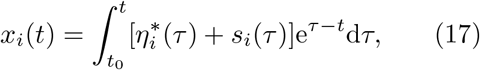

from which the calculation of the dynamical correlation would be straightforward. Specifically, assuming independent the external drive on different units, when *t* − *t*_0_ → ∞ (steady state) we have

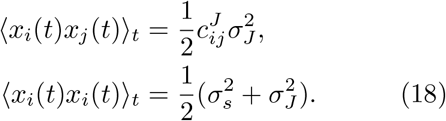

The dynamical correlation 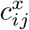 is then directly calculated according to equation (6). As results, while the relationship between coupling and dynamical correlation is linear with unity slope in the absence of external drive, a non-zero *σ*_*s*_ does not affect the linearity, but will suppress the slope.

### 11.2 Numerical simulations of network dynamics

In this study we simulate the dynamics of networks with the coupling matrix elements drawn from Gaussian distributions (Gaussian networks) to verify the theoretical predictions, and further simulated the networks with heavy-tailed coupling distribution and networks with small-world property to see numerically whether the conclusions in the Gaussian networks remain applicable. The simulations can be divided into three steps: first constructing the coupling correlation matrix **C**, then generating random realizations of coupling matrices **J** from a fixed coupling correlation ^)^ matrix, and finally simulating the dynamics.

#### 11.2.1 Constructing coupling correlation matrix

In this work we investigate two types of coupling correlation matrices marked by their distinct spectral property: coupling correlation matrices with bulk spectra, and those with long-tailed spectra.

We construct the coupling correlation matrices with two approaches. In the first one, we begin with generating a Gaussian random matrix of size *N* × *N* . Although the correlations between different rows are expected to vanish theoretically, the entries of sample correlation matrix will randomly fluctuate around zero due to the finite sample effect. Therefore, this sample correlation matrix is fixed as the coupling correlation matrix **C**. This approach is straightforward, but only apply for the coupling correlation matrix with bulk spectrum, because the sample correlation matrix of a Gaussian random matrix have eigenvalues that follow the Marchenko–Pastur distribution that is a bulk distribution, as a classic result of the random matrix theory (**Supplementary Fig.1**) The results of **Fig.2** and **Fig.3a,b** are based on this approach.

The second approach constructs coupling correlation matrix via the spectral decomposition method **C** = **UΛU**^−**1**^, which applies well for both bulk and long-tailed spectra cases. Instead of directly constructing the correlation matrix, we first construct its eigenvalues and eigenvectors. Specifically, the eigenvalues are sampled from a desired spectral distribution, for example uniform, exponential or Poisson for the bulk case, and log-normal or Cauchy for the long-tailed case. Notably, these spectral distributions must have zero probability for negative values, and must have mean equal to one, since the eigenvalues of a correlation matrix must be all positive and sum up to *N* . In the meantime, the eigenvectors **U** is a random orthogonal matrix sampled from the Haar measure on *O*(*N*) (obtained via the QR decomposition of a Gaussian random matrix). This approach allows for more flexible control of the coupling correlation properties by manually designing **Λ** and **U**. An interim correlation matrix **C**_**0**_ is calculated as **C**_**0**_ = **UΛU**^−**1**^. Importantly, since **Λ** and **U** are randomly sampled, the direct multiplication of them does not guarantee the diagonal matrix of **C**_**0**_, denoted as **D**, to be strictly equal to identity. Therefore, the final valid correlation matrix **C** is calculated as **C** = **D**^−**1***/***2**^**C**_**0**_**D**^−**1***/***2**^. This normalization will affect the values of the spectrum, but the influence is mild and will not change the property of spectral distribution (**Supplementary Fig.5**).

#### 11.2.2 Generating the coupling matrix

After constructing the coupling correlation matrix **C**, generating the coupling matrices **J** is straight-forward. Specifically, for each simulation, we generate *N* samples of *N* Gaussian random variables that follow *N* (0, **Σ**) where 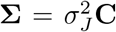. The result is a *N* × *N* coupling matrix with the rows corresponding to variables and columns corresponding to samples, which is the coupling matrix **J**. As definition, the entries of **J** will have standard deviation *σ*_*J*_, with the *i*-th and *j*-th rows being correlated with 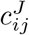.

#### 11.2.3 Simulate network dynamics

With the coupling matrices, we then perform numerical simulations of neural networks according to equation (1), as will be discussed in the **Numerical Estimation of the Largest Lyapunov Exponent** part in detail. For a fixed coupling correlation matrix **C**, we simulate 1000 random realizations of coupling matrices **J**. Therefore, for a given unit pair *i* and *j*, their coupling correlation 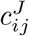 is constant across different network realizations, and their dynamical correlation 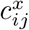 is calculated as the average of the simulated correlation between *x*_*i*_(*t*) and *x*_*j*_(*t*) over all network realizations.

### 11.3 Numerical Estimation of the Largest Lyapunov Exponent

To quantitatively assess the chaoticity of the network dynamics, we numerically compute the largest Lyapunov exponent (LLE), denoted as *λ*_*max*_. The Lyapunov exponent characterizes the average exponential rate of divergence or convergence of nearby trajectories in phase space. A positive LLE is a hallmark of deterministic chaos.

Our computation follow the standard algorithm for continuous-time systems, which involves simultaneously integrating the original nonlinear system along with its linearized equations (the variational equations) for a set of perturbation vectors. In what follows we present the core procedure in the case of zero external drive.

#### 11.3.1 System Integration

The temporal evolution of the network dynamics is numerically integrated using the fourth-order Runge-Kutta method with a time step of Δ*t* = 0.01 and *n*_*T*_ = 10000 time steps. The simulation results about network dynamics are performed at *N* = 500. The system is described by equation (1), with a matrix form representation:

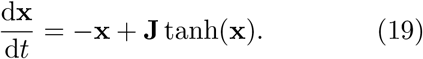

Notice that the nonlinear activation function is taken as *ϕ*(**x**) = tanh(**x**). The initial state **x**(**0**) is randomly sampled from *N* (0, 1).

#### 11.3.2 Tangent Space Evolution

A set of *k* initially orthonormal perturbation vectors {**v**_**1**_, **v**_**2**_, · · ·, **v**_**k**_ } is initialized randomly in the tangent space at **x**(**0**). These vectors represent infinitesimal perturbations to the system’s trajectory. Their evolution is governed by the linearization of the dynamics around the current trajectory **x**(**t**):

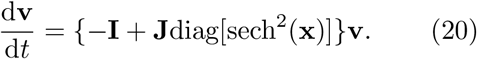

This linearized equation was integrated in parallel with the main system for all *k* vectors using the same time step Δ*t*.

#### 11.3.3 Gram-Schmidt Orthonormalization

To ensure that we accurately track the direction of the fastest growth, which is embedded in the dominant subspace of the linear flow, the set of perturbation vectors was periodically re-orthonormalized every *τ* steps. This was performed using a **QR** decomposition, **V** = **QR**. The orthogonal matrix **Q** provides a fresh set of orthonormal perturbation vectors. By updating **V** = **Q**, we effectively reset the vectors to unit length and ensures their mutual orthogonality, while preserving the dominant subspace they span. This step is crucial for numerical stability, as it prevents the exponential growth of vector magnitudes from causing overflow and maintains a well-conditioned basis for tracking the most unstable directions in the tangent space. The columns of **Q** become the initial conditions for the subsequent iteration of the linearized flow. On the other hand, The upper triangular matrix **R** is used for recording the exponential growth rate. Its diagonal elements, *r*_*ii*_, represent the local growth factors of the perturbation vectors over the preceding orthonormalization interval *τ* . Specifically, |*r*_*ii*_| is the factor by which the *i*-th vector was stretched or compressed within the subspace orthogonal to the first *i*−1 more unstable directions. The natural logarithm of the first element, ln |*r*_11_|, is accumulated at each step. The long-term average of these values yields the largest Lyapunov exponent, isolating the average exponential growth rate along the most unstable manifold, independent of the rotational dynamics captured by the off-diagonal elements of **R**. In essence, the Gram-Schmidt orthonormalization step decouples the measurement of exponential growth (via **R**) from the preparation of vectors for future evolution (via **Q**). This ensures both the accuracy of the exponent calculation and the long-term stability of the numerical algorithm.

#### 11.3.4 Largest Lyapunov Exponent Calculation

Finally, the LLE was estimated from the average growth rate of the first perturbation vector, which, due to the Gram-Schmidt orthonormalization, aligns with the most unstable direction in the tangent space. The estimate is given by:

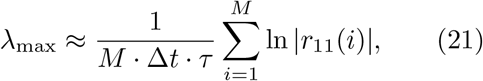

where *r*_11_(*i*) is the first diagonal element of the *R* matrix from the *i*-th orthonormalization step, and *M* is the total number of such steps performed during the simulation.

### 11.4 Numerical simulation of conditions for size-invariant coupling correlation

Although long-tailed spectra can produce coupling correlations of biologically plausible magnitudes, the observed size-invariance of these magnitudes is a non-trivial feature that may require additional conditions to emerge. To examine this, we focus on spectra following a log-normal distribution, which exhibit an appropriate degree of long-tailed nature—being bulk-like on the logarithmic scale—consistent with biological observations.

Using the spectral decomposition approach introduced in the **Numerical simulations of network dynamics** part of **Methods**, we construct coupling correlation matrices while systematically varying two parameters: the network size *N* (ranging from 100 to 3300) and the dispersion parameter of the spectral distribution, *σ*_*λ*_ (ranging from 0.1 to 100). For a given network size *N*, we simulate how the coupling correlation magnitude *σ*_*R*_—averaged over multiple random realizations of the correlation matrix—depends on *σ*_*λ*_. The simulation results yield discrete scatter plots of *σ*_*R*_ as a function of *σ*_*λ*_ (**Supplementary Fig.2a**).

To capture this dependence continuously, we fit the numerical data using a smoothing spline approach (the ‘smoothingspline’ method in MAT-LAB’s `fit` function), obtaining a functional form *σ*_*R*_ = *f* (*σ*_*λ*_; *N*) (**Supplementary Fig.2b**,**c**). This function is monotonically increasing, allowing us to compute its inverse numerically, *σ*_*λ*_ = *f* ^−1^(*σ*_*R*_; *N*), which in turn enables us to solve the inverse problem—namely, determining the value of *σ*_*λ*_ required to achieve a specified level of *σ*_*R*_ (**Supplementary Fig.2d**,**e**).

Finally, by repeating this fitting-and-inversion procedure for different values of *N*, we obtain the relationship between network size and spectral dispersion parameter *σ*_*λ*_ under a fixed target correlation magnitude *σ*_*R*_ (**Supplementary Fig.2f**). This relationship characterizes the conditions under which long-tailed spectra can sustain size-invariant coupling correlation magnitudes.

### 11.5 Biological data Analysis

#### 11.5.1 Open dataset of structural connectivity and neural dynamics

We analyze the publicly available functional and structural datasets of human [26, 27], mouse [28, 31, 32] and *Drosophila* brains [29, 30, 33]. The functional data are recorded by fMRI for human, and wide-field calcium imaging for mouse and *Drosophila*. These data provide the macroscopic neural dynamics in terms of the BOLD (fMRI) or calcium (wide-field imaging) signals of brain regions, which reflect the overall activities of the numerous neurons within the regions. Specifically, the BOLD (blood-oxygen-level-dependent) signal indirectly reflects neural activity through neurovascular coupling, where neuronal firing increases metabolic demand, triggering a hemodynamic response that alters the balance of oxygenated/deoxygenated hemoglobin. Meanwhile, wide-field calcium imaging directly measures neural activity via calcium-sensitive indicators (e.g., GCaMP), which fluoresce in response to action potentials and synaptic activity. These functional data have undergone preliminary preprocessing, for example motion correction, global signal regression, template registration, filtering, etc.

On the other hand, the structural data provide the macroscopic coupling strength between brain regions that reflect the overall connectivity composed of all the axonal projections formed by single neurons. Specifically, for human we use the diffusion tensor imaging (DTI) data that maps white matter structural connectivity between brain regions by modeling water diffusion anisotropy along axonal tracts, offering macroscopic fiber pathway reconstruction. For mouse, the neural bulk tracing techniques inject viral tracers or dyes to a source region, which will then be absorbed by local neuronal population, travel along their axonal projections and enter downstream neurons in target areas to reveal the collective axonal projections from source to target. Finally, for *Drosophila* the single-neuron level synaptic connections were first measured using electron microscope and then combined to generate population level connectivity according to the partitioning of neurons into brain regions.

Specifically, for the first human dataset, structural connectivity matrices and neural dynamics were obtained from the open-access dataset provided by Jimenez-Marin et al [26]. The original data were derived from 136 healthy subjects from the Human Connectome Project (HCP). Brain parcellation was performed using a data-driven approach based on spatially constrained spectral clustering, yielding multiple parcellation scales. The second human dataset provided results derived from the MICA open dataset [27], including parcellation based on Desikan-Killiany atlas, Destrieux atlas, subparcellations within Desikan-Killiany atlas, Glasser atlas and Schaefer atlas.

For further experimental details concerning the collection and preprocessing of functional and structural data, see the respective references.

#### 11.5.2 Coupling correlation and dynamical correlation

Each functional data has the shape of *N* × *T* where *N* is the number of brain regions and *T* is the number of time points. On the other hand, each structural data is a square matrix with the shape *N* × *N*, with the diagonal elements equal to zero. The dynamical correlation 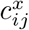 is calculated as the Pearson’s correlation coefficient between of the *i*-th and *j*-th rows in the functional data, while the coupling correlation *ρ*_*ij*_ is calculated as the correlation coefficient between the *i*-th and *j*-th rows in the structural data. At last, the structural coupling matrix and functional dynamical correlation matrix are averaged over subjects (136 subjects for HCP human dataset, 50 subjects for MICA human dataset, 9 mice with in total 29 recording sessions, and 20 *Drosophila*).

#### 11.5.3 Network communication measures

To benchmark our coupling correlation against established approaches, we considered five widely used network communication measures. Let *W* ∈ ℝ^*N* ×*N*^ denote the matrix of structural connectivity weights between *N* brain regions, where *W*_*ij*_ *>* 0 indicates the presence of an anatomical connection between regions *i* and *j*, and *W*_*ij*_ = 0 otherwise. To quantify transmission cost, we define the length matrix *L* = 1*/W*, where *L*_*ij*_ represents the travel distance between *i* and *j*. This remapping from connection weights to lengths is required for communication models that aim to minimize the cost of transmitting signals between regions. The measures were computed using the Brain Connectivity Toolbox and are defined as follows. **Shortest path efficiency** identifies the minimum-cost path between two regions using the Floyd–Warshall algorithm, yielding the minimal transmission cost 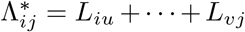 Efficiency is then 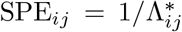. **Navigation efficiency** implements a greedy routing protocol: from source *i*, the signal moves to the neighbor with the shortest Euclidean distance to target *j*, repeating until *j* is reached (success) or a region is revisited (failure). For successful paths, the path length is Λ_*ij*_ = *L*_*iu*_ +· · · + *L*_*vj*_; failed paths are assigned Λ_*ij*_ = ∞. Efficiency is NE_*ij*_ = 1*/*Λ_*ij*_. **Diffusion efficiency** assumes unbiased random walks, with transition matrix *T*_*pq*_ = *W*_*pq*_*/* ∑_*i*_ *W*_*pi*_. Using *T*, one computes ⟨*H*_*ij*_⟩, the expected number of steps for a random walker to travel from *i* to *j*, and defines DE_*ij*_ = 1*/* ⟨*H*_*ij*_⟩ . **Search information** quantifies the information needed to bias a random walker to follow the shortest path. Letting Π_*ij*_ = *T*_*iu*_ *T*_*vj*_ be the probability of traversing the shortest path by chance, search information is SI_*ij*_ = log_2_(Π_*ij*_). **Communicability** sums over all walks between *i* and *j*, weighting each walk inversely by its length: 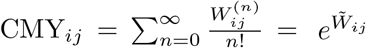, where 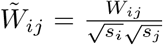 is the normalized connectivity matrix and *s*_*i*_ = ∑_*j*_ *W*_*ij*_ is the strength of region *i*.

## 12 Data availability

The first human dataset (HCP) is available in Zenodo repository at https://zenodo.org/records/8158914 [26], the second human is available at [27]. *Drosophila* data dataset (MICA) https://osf.io/j532r/ are available at https://doi.org/10.6084/m9.figshare.13349282 [29]. The structural data of mouse are available from the Allen Brain Atlas website https://download.alleninstitute.org/publications/ [32], while the functional data are available on ScienceDB (https://www.scidb.cn/en) at https://doi.org/10.57760/sciencedb.17064 [28].

The structural connectivity matrices and dynamical correlation matrices directly used in the main article are attached in the Supplementary Data.

## 13 Code availability

The codes used in this study are attached in the Supplementary Data.

