## Supplementary information for "A Structural Principle for Macroscopic Neural Dynamics Correlations"

### Contents

**1 Details of theoretical analysis** **2**
1.1 Calculation of the generating functional $Z_{\mathbf{J}}$ and averaged generating functional $\bar{Z}$ . . . . . . . . . . . . . . . . . . . . . . . . . . . . . . . . . . 2
1.2 Derivation of the DMFT equations via saddle-point approximation . . 5

**2 The bulk spectrum of the sample correlation matrix of Gaussian random matrix** **7**

**3 Numerical simulation of the condition for size-invariant correlation magnitude** **8**

**4 The relationship between $c^J_{ij}$ and $c^x_{ij}$ under different spectral distributions** **10**

**5 The relationship between $\lambda_i$ and $\sigma_i$ under different spectral distributions** **11**

**6 Normalization of constructed correlation matrix** **13**

### 1 Details of theoretical analysis

As is mentioned in the **Methods** part of the main text, our theoretical analysis can be divided into three steps: 1) calculating the generating functional of the dynamical process with the path integral method, 2) deriving the dynamical mean-field theory equations from the generating functional via saddle-point approximation, and 3) analyzing the dynamical properties from the DMFT equations. Here we present additional details for the previous two steps.

#### 1.1 Calculation of the generating functional $Z_{\mathbf{J}}$ and averaged generating functional $\bar{Z}$

The derivation started from the generating functional $Z_{\mathbf{J}}$ of dynamical process for a given coupling matrix $\mathbf{J}$, which provided a thorough description of the dynamical properties. Then, $\bar{Z}$ is calculated as the averaged generating functional over the random realizations of the coupling matrix. Unlike $Z_{\mathbf{J}}$ that is affected by the specific realization of the coupling matrix, $\bar{Z}$ will reveal the dynamical properties that only depend on the structural statistics of $\mathbf{J}$ (e.g., coupling correlation). Here, unlike the conventional random neural networks that only considered the expectation and variance of the coupling matrix elements, we introduced the coupling correlation that is the seond-order mixed moment that reads $E(J_{ik}J_{jk}) = r^J_{ij}\sigma^2_J$. As the coupling correlation remain invariant across random network realizations, the averaged generating functional $\bar{Z}$ is expected to reveal how this network statistics shapes the dynamics.

We discretized the systems described by equation (1) in the main text while adding the perturbation field $j_i(t)$:

$$\frac{\mathrm{d}x_i(t)}{\mathrm{d}t} = -x_i(t) + \sum_j J_{ij}\phi[x_j(t)] + s_i(t) + j_i(t), \tag{S1}$$

by dividing the time intervals of interest $[t_0, t]$ into $n_T$ segments of length $\delta t$ such that $n_T\delta t = t - t_0$:

$$x_i^{a+1} - x_i^a = -x_i^a\delta t + \sum_{j=1}^{N} J_{ij}\phi_j^a\delta t + s_i^a\delta t + j_i^a\delta t + x_i^0\delta_{a0}^{Kr}. \tag{S2}$$

Here $\phi_j^a$ is the abbreviation of $\phi[x_j(t_a)]$, and $\delta_{a0}^{Kr}$ is the Kronecker delta that sets the initial condition at $t_0$ to be $x_i^0$. The discretization allows for a convenient expression of the probability density of the dynamical path $\{x_i^a\}_{t_a\in[t_0,t]}$ under the constraint of equation (1), which is given by

$$P[x_i^a]_{\mathbf{J}} = \int p(s_i^a)\mathrm{d}s_i^a\delta[x_i^{a+1} - x_i^a + (x_i^a - \sum_j J_{ij}\phi_j^a)\delta t - s_i^a\delta t - j_i^a\delta t - x_i^0\delta_{a0}^{Kr}] \tag{S3}$$

where the Dirac $\delta$ function indicates that the dynamics of a unit $i$ at time $t_a$ takes the probability density of 1 if it satisfies the differential equation, and 0 other wise.

The subscript $\mathbf{J}$ indicates that the probability density is a function of the specific realization of the coupling matrix $\mathbf{J}$. Then, probability density of the whole dynamical path is calculated as the product of different units at different time points, with $P[x]_{\mathbf{J}} = \prod_{i,a} P[x_i^a]_{\mathbf{J}}$. The Fourier representation of the $\delta$-function yields

$$P[x]_{\mathbf{J}} = \int \prod_{i,a} \frac{\mathrm{d}\hat{h}_i^a}{2\pi} \exp\{-\mathrm{i}\hat{x}_i^a[x_i^{a+1} - x_i^a + (x_i^a - \sum_j J_{ij}\phi_j^a)\delta t - \frac{\sigma_s^2}{2}\mathrm{i}\hat{x}_i^a\delta t - j_i^a\delta t - x_i^0\delta_{a0}^{Kr}]\}, \tag{S4}$$

where the conjugate variable $\hat{x}_i$ is naturally introduced. Therefore, the generating functional reads

$$\begin{aligned} Z[j,\hat{j}]_{\mathbf{J}} &= \int \prod_{i,a} \mathrm{d}x_i^a P[x_i^a]_{\mathbf{J}} \exp\{-\mathrm{i}x_i^a\hat{j}_i^a\} \\ &= \int \prod_{i,a} \frac{\mathrm{d}x_i^a \mathrm{d}\hat{x}_i^a}{2\pi} \exp\{-\mathrm{i}\hat{x}_i^a[\frac{x_i^{a+1} - x_i^a}{\delta t} + x_i^a - \sum_j J_{ij}\phi_j^a - \frac{\sigma_s^2}{2}\mathrm{i}\hat{x}_i^a - j_i^a - \frac{x_i^0\delta_{a0}^{Kr}}{\delta t}]\delta t \\ &+ (\mathrm{i}\hat{x}_i^a j_i^a + \mathrm{i}x_i^a\hat{j}_i^a)\delta t\}. \end{aligned}$$

Then, taking the continuum limit $n_T \to \infty$, we got the generating functional $Z[j,\hat{j}]_{\mathbf{J}}$ for continuous systems as the integral over all possible paths of $x_i$ and $\hat{x}_i$ as in equation (2) in the main text:

$$Z[j,\hat{j}]_{\mathbf{J}} = \int \prod_i \mathrm{D}x_i \mathrm{D}\hat{x}_i \exp\{-\sum_i S[x_i,\hat{x}_i] + \sum_{i,a}(\mathrm{i}\hat{x}_i^a j_i^a + \mathrm{i}x_i^a\hat{j}_i^a)\}, \tag{S5}$$

with the functional integral measure defined as $\mathrm{D}x_i \equiv \lim_{n_T\to\infty} \prod_a \mathrm{d}x_i^a$ and $\mathrm{D}\hat{x}_i \equiv \lim_{n_T\to\infty} \prod_a \frac{\mathrm{d}\hat{x}_i^a}{2\pi}$.

The averaged generating functional $\bar{Z}[s,\hat{s}]$ is then calculated as

$$\begin{aligned} \bar{Z}[j,\hat{j}] &= E(Z[j,\hat{j}]_{\mathbf{J}}) \\ &= \int \prod_i \mathrm{D}x_i \mathrm{D}\hat{x}_i E(\exp\{-\sum_i S[x_i,\hat{x}_i]\}) \exp\{\sum_{i,a}(\mathrm{i}\hat{x}_i^a j_i^a + \mathrm{i}x_i^a\hat{j}_i^a)\}. \end{aligned} \tag{S6}$$

Since $S[x_i,\hat{x}_i] = \sum_a \mathrm{i}\hat{x}_i^a(\dot{x}_i^a + x_i^a - \sum_j J_{ij}\phi_j^a - \frac{\sigma_s^2}{2}\mathrm{i}\hat{x}_i^a - x_i^0\delta_{a0})$, the above step is essentially a calculation of the average of $\exp\{\sum_{i,j} J_{ij}\mathrm{i}\hat{x}_i^a\phi_j^a\}$ over random realizations of the coupling matrix $\mathbf{J}$. Given the multivariate Gaussian distribution of $J_{ij}$, the result reads

$$E(\exp\{\sum_{i,j} J_{ij}\mathrm{i}\hat{x}_i^a\phi_j^a\}) = \exp\{\sum_{i,j}\frac{\sigma_J^2}{2}(\sum_a \mathrm{i}\hat{x}_i^a\phi_j^a)^2\}\times$$
$$\times\exp\{\sum_{i\neq j}\sum_k \frac{r_{ij}^J\sigma_J^2}{2}(\sum_a \mathrm{i}\hat{x}_i^a\phi_k^a)(\sum_b \mathrm{i}\hat{x}_j^b\phi_k^b)\}, \tag{S7}$$

By interchanging the order of summation, we got

$$E(\exp\{\sum_{i,j} J_{ij}\mathrm{i}\hat{x}_i^a\phi_j^a\}) = \exp\{\frac{\sigma_J^2}{2}\sum_{a,b}(\sum_i \mathrm{i}\hat{x}_i^a\mathrm{i}\hat{x}_i^b)(\sum_j \phi_j^a\phi_j^b)$$
$$+\frac{\sigma_J^2}{2}\sum_{a,b}(\sum_{i\neq j} r_{ij}^J\mathrm{i}\hat{x}_i^a\mathrm{i}\hat{x}_j^b)(\sum_k \phi_k^a\phi_k^b)\} \tag{S8}$$

.

To simplify the non-local interaction term, we introduced the following change of variables $\hat{\mathbf{x}}^{\mathbf{a}} = \mathbf{Q}\hat{\boldsymbol{y}}^{\boldsymbol{a}}$, $\mathbf{x}^{\mathbf{a}} = \mathbf{Q}\boldsymbol{y}^{\boldsymbol{a}}$, $\hat{\mathbf{j}}^{\mathbf{a}} = \mathbf{Q}\hat{\boldsymbol{l}}^{\boldsymbol{a}}$ and $\mathbf{j}^{\mathbf{a}} = \mathbf{Q}\boldsymbol{l}^{\boldsymbol{a}}$, which leads to

$$\sum_i \mathrm{i}\hat{x}_i^a\mathrm{i}\hat{x}_i^b = \sum_i \mathrm{i}\hat{y}_i^a\mathrm{i}\hat{y}_i^b, \tag{S9}$$

$$\sum_{i\neq j} r_{ij}^J\mathrm{i}\hat{x}_i^a\mathrm{i}\hat{x}_j^b = \sum_i(\lambda_i - 1)\mathrm{i}\hat{y}_i^a\mathrm{i}\hat{y}_i^b, \tag{S10}$$

$$\sum_{i,a}\mathrm{i}\hat{x}_i^a(\dot{x}_i^a + x_i^a - x_i^0\delta_{a0}) = \sum_{i,a}\mathrm{i}\hat{y}_i^a({}_i^a + y_i^a - y_i^0\delta_{a0}), \tag{S11}$$

$$\sum_{i,a}(\mathrm{i}\hat{x}_i^a j_i^a + \mathrm{i}x_i^a\hat{j}_i^a) = \sum_{i,a}(\mathrm{i}\hat{y}_i^a l_i^a + \mathrm{i}y_i^a\hat{l}_i^a), \tag{S12}$$

where the columns of $\mathbf{Q}$ are the eigenvectors of the coupling correlation matrix, and $\lambda_i$ is the $i$-th eigenvalue. The generating functional of the latent system can be calculated from equation (3) in the main text following the above procedure, or alternatively by substituting the latent variables into the expression of $\bar{Z}[j,\hat{j}]$ for the original system. Here, we show the latter approach, substituting equations (S9-S12) into equation (S6) so that the averaged generating functional is expressed as

$$\bar{Z}[l,\hat{l}] = \int |\mathbf{Q}|^2\prod_i \mathrm{D}y_i\mathrm{D}\hat{y}_i \exp[-\sum_{i,a}\mathrm{i}\hat{y}_i^a(\dot{y}_i^a + y_i^a - \frac{\sigma_s^2}{2}\mathrm{i}\hat{y}_i^a - y_i^0\delta_{a0})$$
$$+\frac{\sigma_J^2}{2}\sum_{a,b}(\sum_i \lambda_i\mathrm{i}\hat{y}_i^a\mathrm{i}\hat{y}_j^b)(\sum_j \phi_j^a\phi_j^b)$$
$$+\sum_{i,a}(\mathrm{i}\hat{y}_i^a l_i^a + \mathrm{i}y_i^a\hat{l}_i^a)] \tag{S13}$$

that yields equation (5) in the main text:

$$\bar{Z}[l,\hat{l}] = \int |\mathbf{Q}|^2 \prod_i \mathrm{D}y_i \mathrm{D}\hat{y}_i \exp\{-\sum_i F[y_i,\hat{y}_i] + \sum_{i,a}(\mathrm{i}\hat{y}_i^a l_i^a + \mathrm{i}y_i^a \hat{l}_i^a)\}. \tag{S14}$$

#### 1.2 Derivation of the DMFT equations via saddle-point approximation

We then employed the equality $C^{ab} \equiv \frac{1}{N}\sum_i \phi_i^a \phi_i^b$ to simplify $\bar{Z}[l,\hat{l}]$ by using the Hubbard-Stratonovich transformation

$$1 = \int \mathrm{d}C^{ab}(\mathrm{d}\hat{C}^{ab}/2\pi)\exp\left[-\mathrm{i}\hat{C}^{ab}(C^{ab} - \sum_i \phi_i^a \phi_i^b/N)\right]. \tag{S15}$$

As a result, Eq. 5 is transformed into

$$\bar{Z}[l,\hat{l}] = \int \mathrm{D}C\mathrm{D}\hat{C}\mathrm{e}^{NU[C,\hat{C};l,\hat{l}]}, \tag{S16}$$

with

$$NU[C,\hat{C};l,\hat{l}] = \frac{N}{2}\sum_{ab}\mathrm{i}\hat{C}^{ab}C^{ab} + NV[C,\hat{C};l,\hat{l}], \tag{S17}$$

$$\begin{aligned} NV[C,\hat{C};l,\hat{l}] = \ln\{\int \prod_i \mathrm{D}y_i \mathrm{D}\hat{y}_i \mathrm{e}^{\sum_i W[y_i,\hat{y}_i;C,\hat{C}]} \times \\ \times \mathrm{e}^{\sum_i \sum_a (\mathrm{i}\hat{y}_i^a l_i^a + \mathrm{i}y_i^a \hat{l}_i^a)}\}, \end{aligned} \tag{S18}$$

and

$$\begin{aligned} W[y_i,\hat{y}_i;C,\hat{C}] = \sum_a \mathrm{i}\hat{y}_i^a(\dot{y}_i^a + y_i^a - \frac{\sigma_s^2}{2}\mathrm{i}\hat{y}_i^a - y_i^0\delta_{a0}) \\ - \frac{1}{2}\sum_{a,b}[\mathrm{i}\hat{C}^{ab}\phi_i^a\phi_i^b + \lambda_i C^{ab}\mathrm{i}\hat{y}_i^a \mathrm{i}\hat{y}_i^b] \end{aligned} \tag{S19}$$

In the thermodynamic limit ($N \gg 1$), the saddle-point approximation ensures that

$$\bar{Z}[l,\hat{l}] \approx \bar{Z}_0[l,\hat{l}] = \mathrm{e}^{NU[C_0,\hat{C}_0;l,\hat{l}]} \tag{S20}$$

due to the exponential decay of the value of $\mathrm{e}^{NU[C,\hat{C};l,\hat{l}]}$ away from $\mathrm{e}^{NU[C_0,\hat{C}_0;l,\hat{l}]}$ magnified by large $N$, where $U[C_0,\hat{C}_0;l,\hat{l}]$ in the exponent is the value of $U$ at the point $C_0^{ab} = 1/N\sum_i\langle\phi_i^a\phi_i^b\rangle, \hat{C}_0^{ab} = 1/N\sum_i\langle\mathrm{i}\hat{y}_i^a\mathrm{i}\hat{y}_i^b\rangle$ that satisfies

$$\frac{\partial}{\partial C}U[C_0,\hat{C}_0;l,\hat{l}]|_{C=C_0,\hat{C}=\hat{C}_0} = 0,$$

$$\frac{\partial}{\partial \hat{C}} U[C_0, \hat{C}_0; l, \hat{l}]|_{C=C_0, \hat{C}=\hat{C}_0} = 0. \tag{S21}$$

Due to the normalization condition $\bar{Z}[l, \hat{l} = 0] = \bar{Z}[j, \hat{j} = 0] = 1$, and the correlation functions that involve only $\hat{y}$ can be calculated as the derivative of $\bar{Z}[l, 0]$ over $l$, $\hat{C}_0^{ab} = 1/N \sum_i \langle \mathrm{i}\hat{y}_i^a \mathrm{i}\hat{y}_i^b \rangle_0$ must vanish to 0. Therefore, we have the value of $NU[C_0, \hat{C}_0; l, \hat{l}]$ as

$$NU[C_0, \hat{C}_0; l, \hat{l}] = \ln \{ \prod_i \int \mathrm{D}y_i \mathrm{D}\hat{y}_i \mathrm{e}^{W[y_i, \hat{y}_i; C_0, 0] + \sum_a (\mathrm{i}\hat{y}_i^a l_i^a + \mathrm{i}y_i^a \hat{l}_i^a)} \} \tag{S22}$$

$\bar{Z}_0[l, \hat{l}]$, as the approximation of the averaged generating functional of the dynamics of the whole system, can be subsequently viewed as the product of $N$ individual generating functionals of single-site dynamics:

$$\bar{Z}_0[l, \hat{l}] = \prod_i Z_i[l, \hat{l}], \tag{S23}$$

where

$$Z_i[l, \hat{l}] = \int \mathrm{D}y_i \mathrm{D}\hat{y}_i \mathrm{e}^{W[y_i, \hat{y}_i; C_0, 0] + \sum_a (\mathrm{i}\hat{y}_i^a l_i^a + \mathrm{i}y_i^a \hat{l}_i^a)} \tag{S24}$$

.

Based on the equality

$$\exp [\sum_{a,b} \lambda_i C_0^{ab} \mathrm{i}\hat{y}_i^a \mathrm{i}\hat{y}_i^b] = \langle \exp [\sum_a \lambda_i^{1/2} \mathrm{i}\hat{y}_i^a \gamma^{*a}] \rangle_t \tag{S25}$$

where $\gamma^*$ is a Gaussian white noise with $\langle \gamma^{*a} \gamma^{*b} \rangle_t = g^2 C_0^{ab}$, we notice that

$$Z_i[l, \hat{l}] = \langle \int \mathrm{D}y_i \mathrm{D}\hat{y}_i \mathrm{e}^{\sum_a \mathrm{i}\hat{y}_i^a (\dot{y}_i^a + y_i^a - \lambda_i^{1/2} \gamma^{*a} - \frac{\sigma_s^2}{2} \mathrm{i}\hat{y}_i^a - y_i^0 \delta_{a0})} \times \\ \times \mathrm{e}^{\sum_a (\mathrm{i}\hat{y}_i^a l_i^a + \mathrm{i}y_i^a \hat{l}_i^a)} \rangle_{\varepsilon_i}. \tag{S26}$$

This indicates $Z_i[l, \hat{l}]$ is the generating functional of the stochastic process

$$\frac{\mathrm{d}y_i}{\mathrm{d}t} = -y_i(t) + \hat{s}_i(t) + \lambda_i^{1/2} \gamma^*(t) + l_i(t), \tag{S27}$$

where $\hat{s}_i(t)$ is the external drive, $\lambda_i^{1/2} \gamma^*(t)$ is the mean-field term representing the internal interactions, and $l_i(t)$ is the perturbation field used for DMFT derivation.

Finally, a linear transformation of the mean-field equations for latent dynamics yields the DMFT equations for the original dynamics $x$ as equation (7) in the main text:

$$\frac{\mathrm{d}x_i}{\mathrm{d}t} = -x_i + s_i(t) + \eta_i^*(t) + j_i(t) \tag{S28}$$

with the Gaussian white noise field $\eta_i^* = \sum_k Q_{ik} \lambda_k^{1/2} \gamma_k^*$. The perturbation field $j_i(t)$ is then taken as zero in the following part.

#### 2 The bulk spectrum of the sample correlation matrix of Gaussian random matrix

When simulating the dynamics of correlated random neural networks in the main analysis, we construct the coupling correlation matrix as the sample correlation of a Gaussian random matrix. In random matrix theory, such a matrix corresponds to a (normalized) Wishart matrix, whose eigenvalue spectrum follows the Marchenko–Pastur distribution in the large-$N$ limit. The Marchenko–Pastur distribution has a characteristic bounded support, with the eigenvalue density being nonzero only within a finite interval $[\lambda_-, \lambda_+]$ and vanishing outside this range. The probability density $f(x)$ of the eigenvalue reads

$$f(x) = \frac{\sqrt{(\lambda_+ - x)(x - \lambda_-)}}{2\pi\sigma_J^2 \lambda x} \tag{S29}$$

for $\lambda_- \leq x \leq \lambda_+$, and $f(x) = 0$ otherwise. When $\mathbf{C}$ is the sample correlation matrix of $N \times N$ square matrix $\mathbf{J}$, we have $\lambda = \frac{N}{N} = 1$ with $\lambda_\pm = \sigma_J^2(1 \pm \sqrt{\lambda})^2$ (**Supplementary Fig.1**).

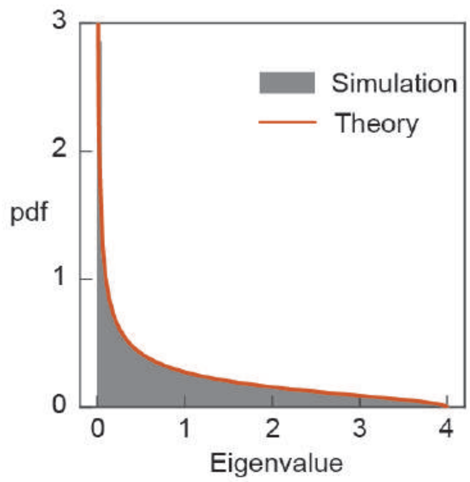


**Supplementary Fig. 1 The bulk spectral distribution of the sample correlation matrix of Gaussian random matrix.** The gray histogram is the distribution of eigenvalues of simulated sample correlation matrices, and the red line is the theoretical Marchenko-Pastur distribution. Notably, the distribution is bulk-like, with eigenvalues bounded between $\lambda_- = 0$ and $\lambda_+ = 4$ ($\sigma_J = 1$)

.

#### 3 Numerical simulation of the condition for size-invariant correlation magnitude

We consider the spectra that follow Log-normal distribution controlled by a dispersion parameter $\sigma_\lambda$. The detailed simulation procedure is provided in **Methods**, and is visualized here step-by-step to enhance clarity. The goal is to learn how the dispersion parameter $\sigma_\lambda$ must scale with system size $N$ to ensure coupling correlation scale $\sigma_R$ remains size-invariant (**Supplementary Fig.2**).

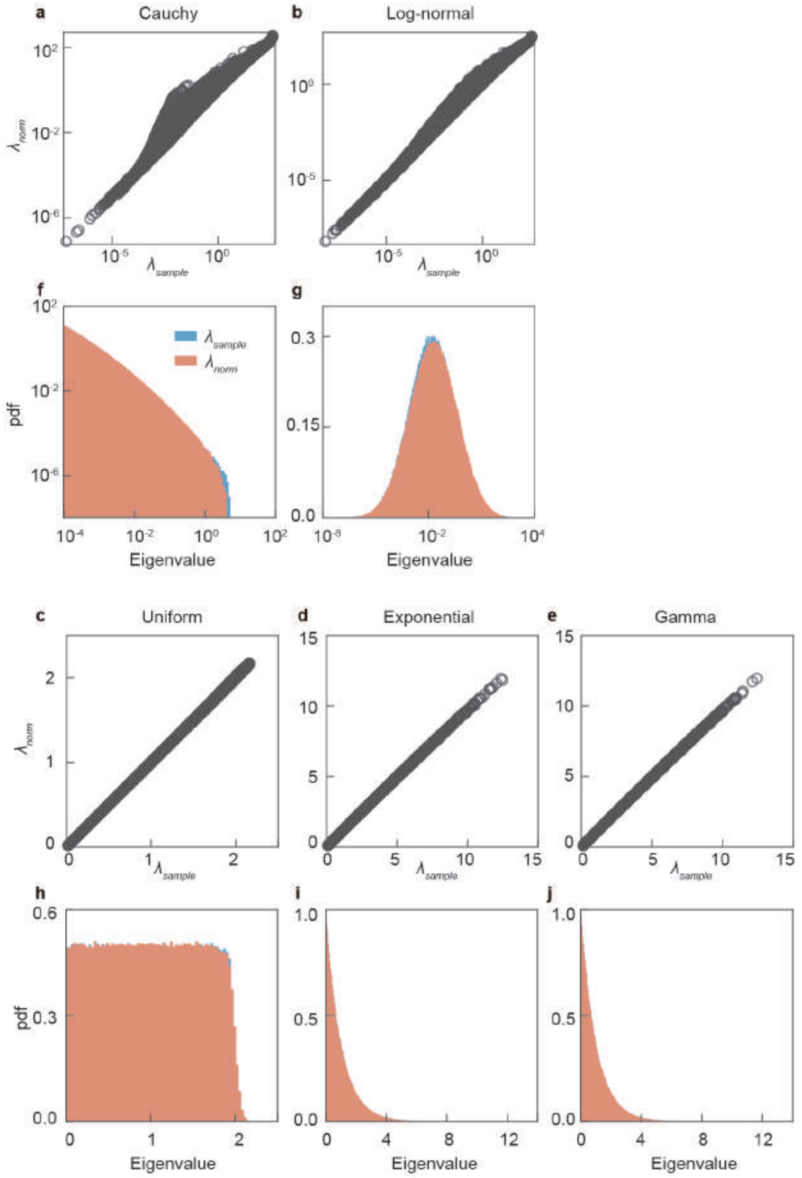


**Supplementary Fig. 2 Numerical simulation of the condition for size-invariant correlation magnitude.** **a**, Dependence of the coupling correlation scale $\sigma_R$ on the dispersion parameter $\sigma_\lambda$, with darker colors indicating larger system sizes (ranging from $N = 100$ to 3301). Each point represents the mean $\sigma_R^2$ averaged over multiple random coupling matrix realizations. **b**, Smoothing spline fits (lines) to the $\sigma_R$–$\sigma_\lambda$ relationship for each $N$ (circles with the same color). **c**, A detailed view confirming the accuracy of the fits. **d**, Inverse mapping from a target $\sigma_R = 0.3$ (dashed line, corresponding to $\mathbb{E}[\sigma_R^2] = 0.09$) to the required $\sigma_\lambda$ for each $N$. **e**, Close-up of the intersections (red crosses) indicating the $\sigma_\lambda$ value for each system size. **f**, The resulting scaling of $\sigma_\lambda$ with $N$.

.

### 4 The relationship between $c_{ij}^J$ and $c_{ij}^x$ under different spectral distributions

Our numerical simulations reveal a consistent relationship between coupling and dynamical correlation that is largely independent of the specific spectral distribution: it remains precisely linear under bulk-like distributions and approximately linear under long-tailed ones. This result not only validates the effectiveness of DMFT for bulk distributions but also indicates that a common origin may be responsible for its failure in long-tailed cases (**Supplementary Fig.3**).

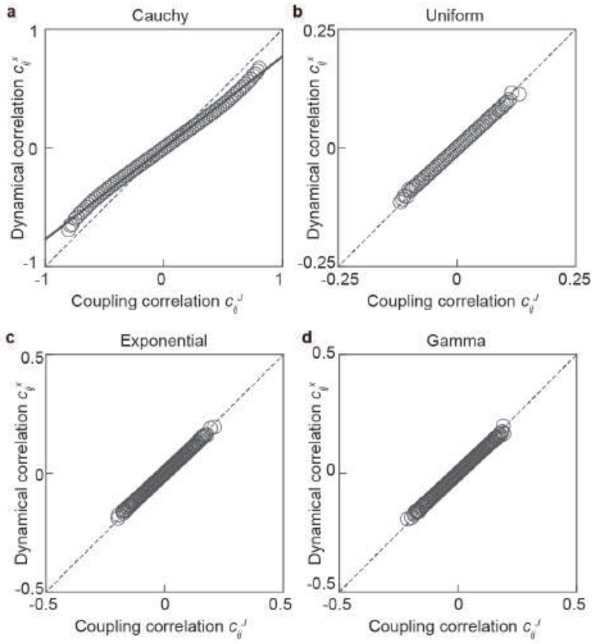


**Supplementary Fig. 3 The relationship between $c_{ij}^J$ and $c_{ij}^x$ under different spectral distributions. a-d**, The simulated relationship when the spectral distribution are Cauchy, Uniform, Exponential and Gamma, respectively. The Cauchy case closely matches the Log-normal results reported in the main text. On the other hand, the other three distributions exhibit a consistently linear relationship between $c_{ij}^J$ and $c_{ij}^x$, despite variations in the magnitude of coupling correlation.

.

#### 5 The relationship between $\boldsymbol{\lambda_i}$ and $\boldsymbol{\sigma_i}$ under different spectral distributions

A key premise for establishing DMFT is that the latent dynamics $y_i(t)$ experience homogeneous collective interaction $\gamma_i(t)$. This premise can be tested by examining whether the latent dynamics scale $\sigma_i$ scales proportionally with the square root of corresponding eigenvalue $\lambda_i$. Simulations confirm that this proportionality holds for bulk-like spectral distributions but breaks down under long-tailed ones, thereby providing a unified explanation for the domain of DMFT's validity (**Supplementary Fig.4**).

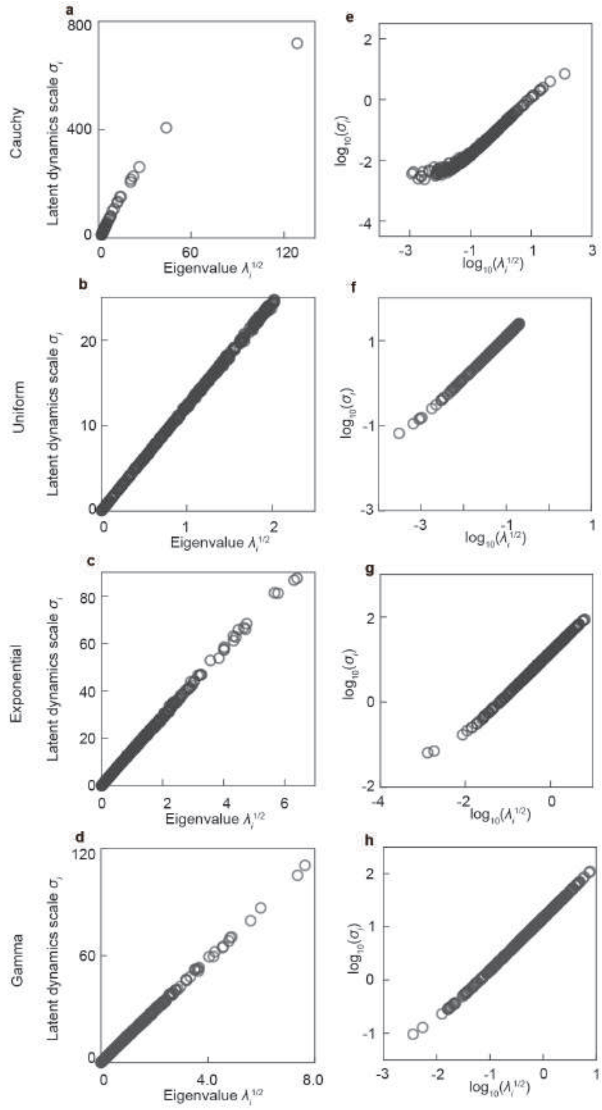


**Supplementary Fig. 4 The relationship between $\lambda_i$ and $\sigma_i$ under different spectral distributions.** The left panels (**a–d**) show the relationship between the latent dynamics scale $\sigma_i$ ($y$-axis) and the corresponding eigenvalue square root $\lambda_i^{1/2}$ ($x$-axis) for the Cauchy, uniform, exponential, and gamma distributions, respectively. The right panels (**e–h**) present the same relationships on a log–log scale.

.

### 6 Normalization of constructed correlation matrix

When constructing the coupling correlation matrix via the spectral decomposition approach, we apply a normalization step $\mathbf{C} = \mathbf{D}^{-1/2}\mathbf{C_0}\mathbf{D}^{-1/2}$, where $\mathbf{D} = \text{diag}(\mathbf{C_0})$, and $\mathbf{C_0} = \mathbf{U}^{-1}\mathbf{\Lambda}\mathbf{U}$ is formed by directly combining randomly generated eigenvectors $\mathbf{U}$ and eigenvalues $\mathbf{\Lambda}$. Here, we demonstrate that the applied normalization has negligible effect on the spectral density, as the eigenvalue probability density function remains fundamentally unaltered throughout the transformation (**Supplementary Fig.5**).

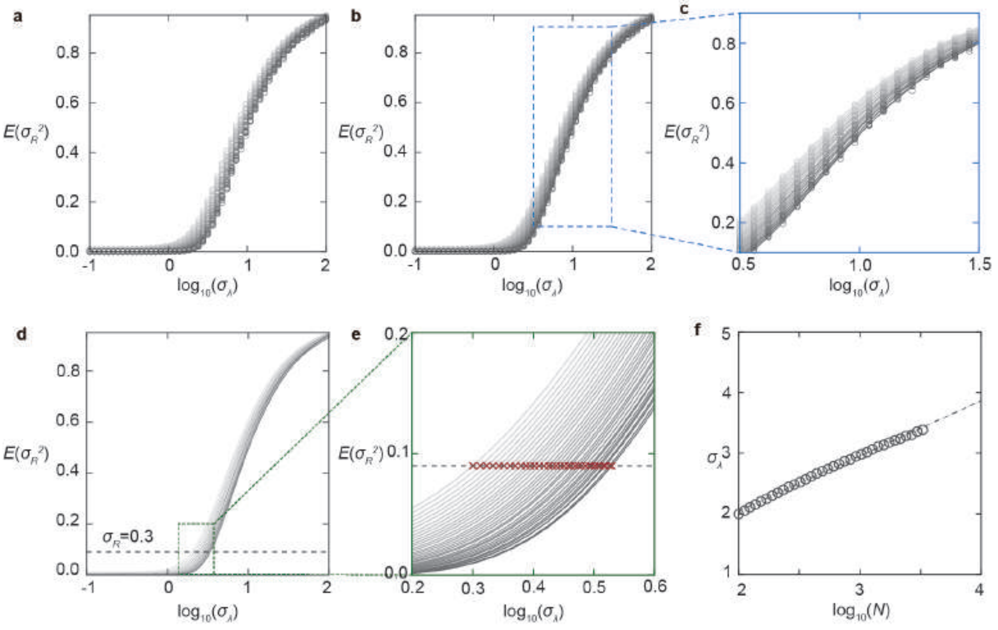


**Supplementary Fig. 5 The spectral distribution is conserved after normalization of coupling correlation matrix.** The upper panels (**a**–**e**) compare the initially sampled eigenvalues ($\lambda_{\text{sample}}$) against their normalized counterparts ($\lambda_{\text{norm}}$) for Cauchy, log-normal, uniform, exponential, and gamma spectral distributions, respectively. Although $\lambda_{\text{norm}}$ occasionally deviates from $\lambda_{\text{sample}}$, particularly in the long-tailed cases (**a,b**), their relationship is well-approximated by the line $y = x$. The corresponding lower panels (**f**–**j**) display the probability distributions of both $\lambda_{\text{sample}}$ and $\lambda_{\text{norm}}$.

.
